# Accurate predictions of conformational ensembles of disordered proteins with STARLING

**DOI:** 10.1101/2025.02.14.638373

**Authors:** Borna Novak, Jeffrey M. Lotthammer, Ryan J. Emenecker, Alex S. Holehouse

## Abstract

Intrinsically disordered proteins and regions (collectively IDRs) are found across all kingdoms of life and play critical roles in virtually every eukaryotic cellular process. In contrast to folded proteins, IDRs lack a stable 3D structure and are instead described in terms of a conformational ensemble, a collection of energetically accessible interconverting structures. This unique structural plasticity facilitates diverse molecular recognition and function; thus, a convenient way to view IDRs is through their ensembles. Here, we combine advances in physics-based force fields for IDPs with the power of modern multi-scale generative modeling to develop STARLING, an approach for the rapid and accurate prediction of IDR ensembles directly from sequence. STARLING enables ensembles of hundreds of conformers to be generated in seconds and works on GPUs and CPUs. This, in turn, dramatically lowers the barrier to the computational interrogation of IDR function through the lens of emergent biophysical properties complementing bioinformatic protein sequence analysis. We evaluate STARLING’s accuracy against extant experimental data and offer a series of vignettes illustrating how STARLING can enable rapid hypothesis generation for IDR function and aid the interpretation of experimental data.

## INTRODUCTION

Intrinsically disordered proteins and protein regions (IDRs) are structurally heterogeneous protein regions that are estimated to constitute ∼30% of eukaryotic proteomes^1,2^. Despite lacking a fixed structure, IDRs play key roles in essential cellular processes such as transcription, translation, and cell signaling^1,3^. Due to their structural heterogeneity, IDRs are best described by a conformational ensemble - a dynamic set of rapidly interchanging conformations^1,4^. Just as folded domains follow a sequence-structure relationship, IDRs follow an analogous sequence-ensemble relationship, where the underlying sequence chemistry dictates the conformations that make up their ensembles^5,6^. IDR ensembles can play essential roles in IDR function and may be perturbed in disease^7–9^. Consequently, just as structural biology has been instrumental in understanding the molecular basis for folded domain function, there is a growing appreciation that the characterization of IDR ensembles is important for understanding IDR function.

Experimental techniques commonly used to interrogate sequence-ensemble relationships include single-molecule Förster Resonance Energy Transfer (smFRET) spectroscopy^10^, Nuclear Magnetic Resonance (NMR) spectroscopy^11^, and Small-Angle X-ray Scattering (SAXS)^12^. While these report on specific aspects of IDR ensembles, they fall short of providing a holistic description of the distribution of conformers (referred to here as a ‘full structural ensemble’). Integrating all-atom simulations with experimental data has proven one effective route to construct full structural ensembles^9,13–17^. However, these simulations require deep technical expertise to ensure reliable conclusions are drawn and are computationally expensive, often requiring significant time to achieve high quality data. While recent advances in coarse-grained (CG) simulations^18,19^ offer a faster alternative, even CG simulations can still take hours to obtain sufficient sampling, making them inappropriate for large-scale investigation into sequence-ensemble relationships. Moreover, CG simulations still require a relatively high level of technical expertise to setup, run, and analyze. Recent deep learning predictors trained on CG simulations have enabled proteome-scale predictions for average values for some subset of observables (e.g. end-to-end distance) but are limited by which observables are predictable^20,21^.

Deep learning approaches such as AlphaFold^22,23^, RosettaFold^24,25^, ESMFold^26,27^, and others^28^ have transformed the field of protein structure predictions, significantly reducing the barrier to the large-scale exploration of sequence-structure relationships (**Fig. 1A**). However, these methods are poorly suited to investigate IDRs^29^ owing to a reduction in alignment-based conservation in disordered proteins, a paucity of appropriate experimental training data, and their training objective (predicting a single best structure from sequence, whereas IDRs should be described by large ensembles of structurally heterogeneous conformers) (**Fig. 1C)**. In short, while we now possess easy-to-use tools for accurately predicting the 3D structure of individual folded domains, equivalent tools for predicting structural ensembles of IDRs are lacking.

**Figure 1.**
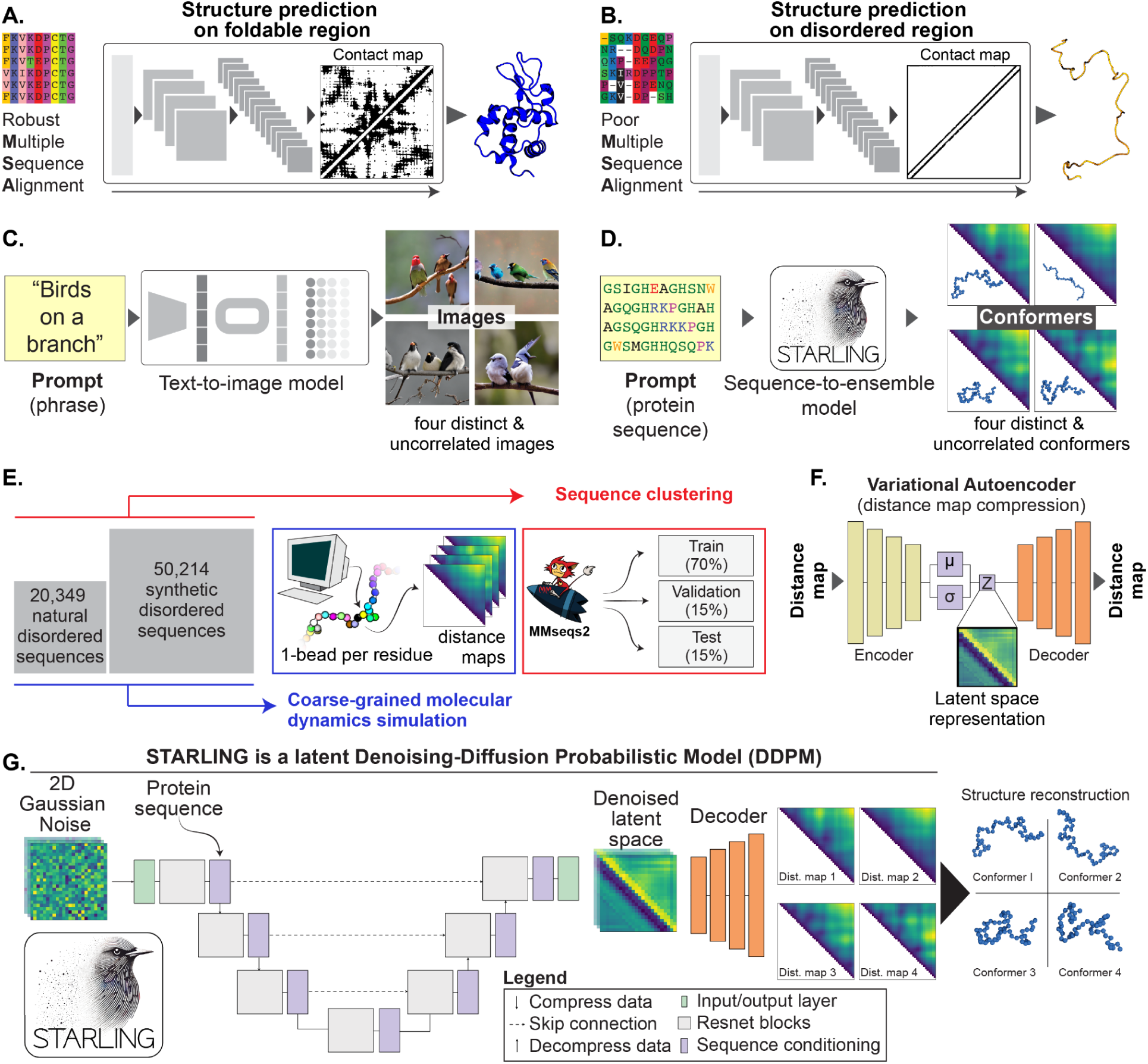
**(A)** Deep learning has revolutionized protein structure prediction, with major advances being facilitated by large-scale evolutionary information. **(B)** Structure prediction methods for folded domains are poorly suited for predicting the behavior of IDRs. These limitations come from the absence of a native-state structure and because evolutionary information is often poorly captured in MSAs. **(C)** Generative text-to-image models enable the creation of many unique and independent images consistent with a single input prompt. **(D)** IDR ensemble generation shares many similarities with text-to-image generation; we require many distinct and uncorrelated conformers, all of which are consistent with an input prompt (amino acid sequence). **(E)**. STARLING is trained on 70,563 amino acid sequences. For each sequence, hundreds of distinct conformers are generated using coarse-grained molecular dynamics (MD) simulations, and each conformer is converted into a distance map (an image). Sequences are separated for training, testing, and validation. **(F)** STARLING makes use of a variational autoencoder (VAE) to compress distance maps to a latent space, allowing a denoising diffusion model to work in this latent space (‘latent diffusion’). **(G)** The overall architecture of the STARLING model combines a latent-space probabilistic denoising diffusion model with a U-Net architecture using residual convolutions and transformer blocks. Latent space maps are decoded to real-space via the VAE decoder. Finally, if needed, distance maps are reconstructed into 3D coordinates via a parallelized multidimensional scaling approach.

Here, we address these challenges by developing a fast and accurate approach for predicting full coarse-grained disordered protein structural ensembles directly from the amino acid sequence. Our approach leverages advances in generative modeling, a deep learning technique capable of creating new and original data. However, developing a generative model poses a key challenge: the need for large training datasets. To address this, we performed large-scale coarse-grained simulations to generate full structural ensembles across tens of thousands of natural and synthetic IDRs. The resulting predictor, **conSTruction of intrinsicAlly disoRdered proteins ensembles efficientLy vIa multi-dimeNsional Generative models** (STARLING) furthers our descent into acronym madness but also allows us to generate full structural ensembles directly from sequence in seconds. A major goal in developing STARLING was to avoid hardware barriers. While STARLING is fast (∼25 conformers/second) on GPUs, it can still generate ensembles in minutes on Intel/AMD CPUs and seconds on Apple CPUs.

STARLING-generated ensembles show good agreement with experimental data, enabling *de novo* exploration of uncharacterized IDRs or aiding in the biophysical interpretation of experimental data. Moreover, here we show proof-of-concept that STARLING can be used to (i) investigate sequence-ensemble relationships for disordered proteins, (ii) explore bound-state conformational ensembles for binary disordered protein complexes, (iii) facilitate rational design of disordered proteins with specific ensembles. Taken together, we propose that STARLING’s ease of use and speed make it a powerful tool for democratizing large-scale exploration of sequence-ensemble relationships in IDRs.

## METHODS & RESULTS

Generative artificial intelligence (AI) has been transformative for text-to-image generation^30–35^. In text-to-image generative AI, an image is generated by passing a prompt (a short phrase describing the desired image) to a pre-trained deep-learning model. That model then generates an image consistent with the prompt, a process referred to as inference. Deep learning models capable of inference must first be trained. For models used in modern text-to-image generative AI, the training is not simply memorizing the images seen in training but learning the relationship between prompts and the features within the associated images. As a result, once a model is trained, if the same prompt is reused many times, it will generate many independent images entirely distinct from any of the individual instances observed in training. Notably, despite being different from one another, every image generated should be consistent with the prompt (**Fig. 1C**).

The mapping of a single text prompt to a collection of different images – each of which is consistent with the input prompt – is precisely the problem we wish to solve for IDR ensemble generation. In IDR ensemble generation, we want to take a text prompt (amino acid sequence) and generate a collection of many distinct and uncorrelated IDR conformations consistent with that prompt (**Fig. 1D**). Moreover, we want this generation process to be fast (seconds) and possible on commodity hardware (laptops and desktops). To achieve this goal, we combine a variational autoencoder (VAE) with a discrete-time denoising-diffusion probabilistic model (DDPM) to create a latent diffusion model^31^. The resulting method (STARLING) enables the accurate and rapid prediction of coarse-grained conformational ensembles of IDRs.

STARLING is trained on coarse-grained simulations of 70,563 rationally designed and naturally occurring disordered protein sequences ranging between **10** and **384** residues in length (**Fig. 1E**, **Fig. S1**). Rationally designed sequences were designed using GOOSE^36^. While naturally occurring IDRs provide training data focused on the manifold of sequence features we may care about the most, the inclusion of a large number of rationally designed sequences allows us to construct training data that systematically titrates across sequence space to provide a well-rounded training set capturing the extremes of sequence composition and patterning. In short, synthetic sequences offer a well-rounded training set, capturing the extremes of sequence space that may not be well represented in natural sequences.

A necessary decision in deep learning models is defining the limits of your training data. We focused on sequences up to 384 amino acids in length for several reasons. First, almost 95% of naturally occurring IDRs across common model organisms are shorter than 384 amino acids (**Fig. S2**). Second, this length enables well-sampled coarse-grained simulations to be completed on a reasonable timescale. Third, the underlying architecture of the models requires us to have a fixed upper limit (see *Methods*). Lastly, experimental characterization of disordered protein sequences longer than ∼350 residues is lacking, making it difficult to assess the validity of model predictions relative to experiments for longer lengths.

Molecular Dynamics (MD) simulations were performed using the Mpipi-GG forcefield. Mpipi-GG^20^ – a variant of the original Mpipi forcefield^18^ – is a one-bead-per-residue coarse-grained model developed for disordered regions. Once run, conformations from simulations of a sequence of length *n* are converted into distance maps, *n* x *n* matrices where each element describes the distance between the i^th^ and j^th^ residue for a specific conformation. This converts each IDR conformation into an “image”, allowing us to leverage innovations developed for the conditional generation of images directly. Overall, our training dataset comprised nearly 10 million distance maps.

Two core limitations of DDPM models are their significant memory needs and a slow generative process at inference. These limitations come from performing the reverse denoising process in complex and high-dimensional data (e.g., images of size 384 x 384 pixels). To mitigate this, we developed a variational autoencoder (VAE) to compress each distance map into a lower-resolution latent space (24 x 24) (**Fig. 1F**). The denoising diffusion process can then occur in this latent space (latent diffusion), significantly reducing memory requirements and inference time. Given this, STARLING was trained in two independent stages.

We first trained a highly accurate VAE that enables the compression of full-resolution distance maps into latent space (**Fig. S3**). Our VAE uses a ResNet18 architecture and learns parameters for an encoder (full-resolution to latent space) and a decoder (latent space to full resolution). We assessed the VAE’s accuracy by encoding and decoding distance maps derived from sequences substantially different from those used in the training and validation sets. We evaluated our model on a held-out test set comprising 10,437 sequences, totaling 1,425,849 distance maps. The model achieved a root mean squared reconstruction error of 1.31 Å (**Fig. S4**). Furthermore, our model accurately reconstructs bond lengths with a root mean squared reconstruction error of 0.19 Å (**Fig. S4**), crucial for accurately modeling protein conformations.

In the second stage of training, we developed a discrete-time denoising-diffusion probabilistic model (DDPM) (**Fig 1D, Fig. S5**). Briefly, this model was trained to learn model parameters that map from random noise to individual latent-space conformer distance maps conditioned on the associated amino acid sequence. The training data came from 9,621,455 distance maps taken from 70,563 simulations. Each map was first compressed to latent space, and then the fixed forward diffusion process was used to add noise to each latent-space distance map. The model was then trained to learn parameters that reverse this forward diffusion process conditioned on the input amino acid sequence (**Fig. 1G**). Ultimately, ensemble generation is enabled by running many rounds of inference in parallel, leading to large numbers of independent distance maps.

While training requires the VAE encoder to generate latent space distance maps, only the VAE decoder is required for inference once the model is trained. Ultimately, the fully trained model (STARLING) combines two different models (the VAE decoder and the DDPM) that work together to enable rapid ensemble prediction. Using default settings, STARLING can generate 200 independent IDR conformations in approximately 9 seconds on a GPU (Nvidia A4500), 30 seconds on a Macbook Pro M3 CPU, and ∼8 minutes on an Intel CPU (Intel(R) Xeon(R) Silver 4210R CPU @ 2.40GHz) (**Figs. S6-8**). Importantly, prediction runtime and memory do not scale with sequence length; a 50 amino acid IDR generates in the same number of seconds with the same memory requirements as a 350 residue IDR (**Fig. S7B**).

We first checked that the STARLING-derived ensemble-averaged global dimensions agree with those obtained from the Mpipi-GG simulation of unseen sequences. In all cases, STARLING ensembles comprised 1000 conformers. Using a held-out test set of 10,437 sequences, we performed Mpipi-GG simulations and calculated radii of gyration (R_g_s) and end-to-end distances (R_e_s). Our STARLING-derived R_g_s (RMSE = 0.78 A, R^2^ = 0.997) and R_e_s (RMSE = 3.21, R^2^ = 0.991) are in excellent agreement with the simulations (**Fig. 2A,B**).

**Figure 2.**
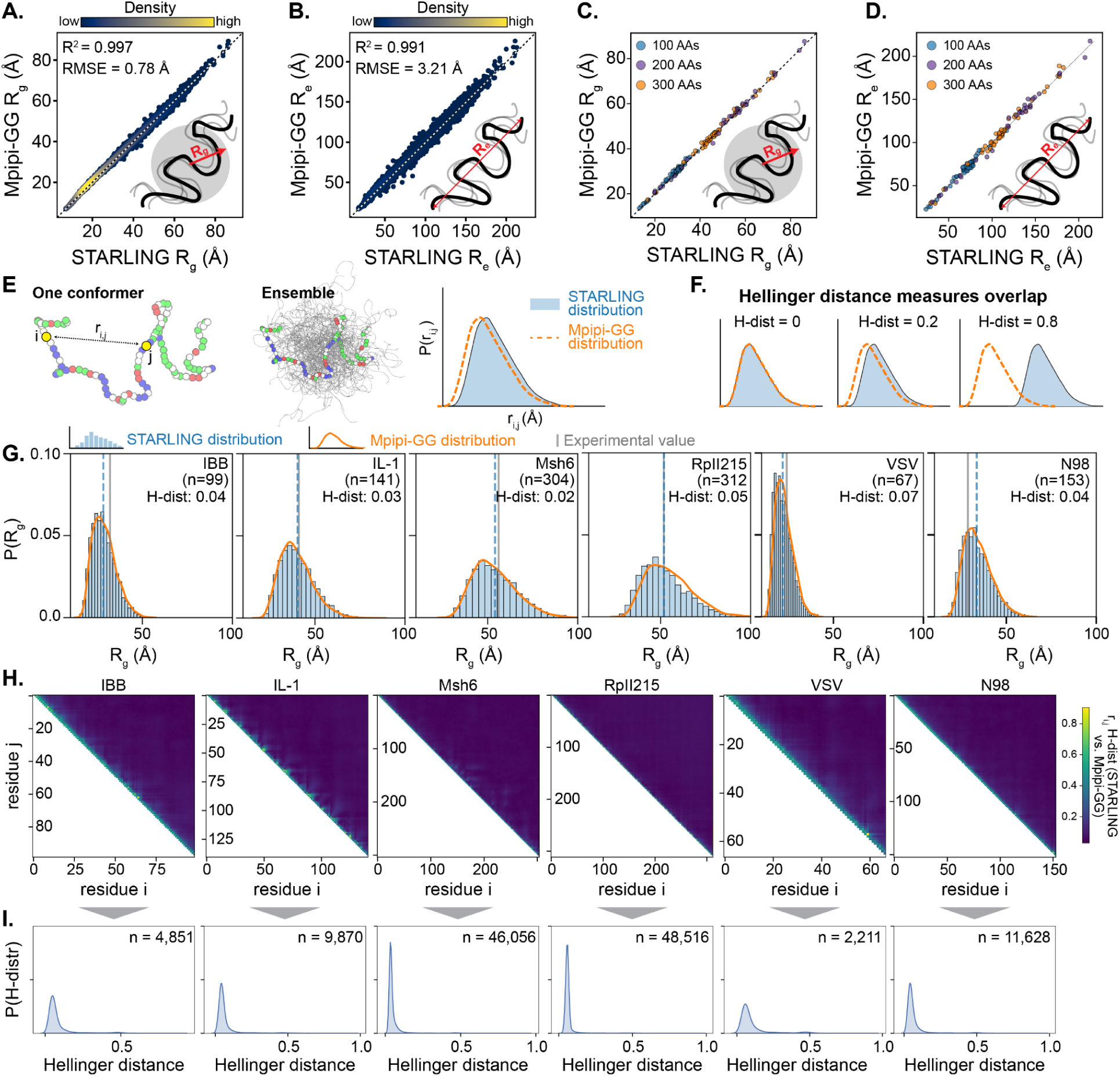
**(A)** Comparison of STARLING-derived average radii of gyration (R_g_s) with those from Mpipi-GG (simulations) from a set of 10,437 unseen sequences. **(B)** Comparison of STARLING-derived average end-to-end distances (R_e_s) with those from Mpipi-GG (simulations) from 10,437 unseen sequences. **(C)** Comparison of STARLING-derived average radii of gyration (R_g_s) for length-matched sequences with a broad range of chemistries for those from Mpipi-GG (simulations) from a set of 137 unseen sequences. This is done to verify STARLING has learned real sequence chemistry, not just that longer sequences are bigger. **(d)** Comparison of STARLING-derived average radii of gyration (R_e_s) for length-matched sequences with a broad range of chemistries for those from Mpipi-GG (simulations) from a set of 137 unseen sequences. This is done to verify STARLING has learned real sequence chemistry, not just that longer sequences are bigger. **(E)** STARLING generates full conformational ensembles, enabling ensemble distributions to be calculated (e.g., R_g_, R_e_, or r_i,j_, inter-residue distance). For example, each (i,j) pair of residues in an ensemble yields a distribution of inter-residue distances, which can be quantified in STARLING ensembles and Mpipi-GG simulations. **(G)** Distribution similarity can be quantified by the Hellinger distance (H-dist), where H-dist = 0 is perfect overlap and H-dist = 1 is no overlap. **(G)** Overlap of radii of gyration distributions obtained from STARLING (blue bars) and Mpipi-GG (orange line) with the experimental value shown for completeness. Overlap between the STARLING and Mpipi-GG distributions is quantified by the Hellinger distance (H-dist). **(H)** Hellinger distance values for all possible i,j inter-residue distance distributions across the same sequences examined in panel E. Almost all i,j distributions overlap very well, with deviations happening only at very short-range distances where the absolute magnitude of poor overlap is sub-angstrom. **(I)** Histogram of Hellinger distances in panels H.

An important determinant of model accuracy reflects ensemble size and the number of denoising steps. Using long timescale Mpipi-GG simulations to obtain an estimate of the true mean for the R_g_, we systematically investigated how ensemble size influences the difference between the STARLING ensemble average value and the Mpipi-GG derived estimated mean (**Fig. S9**). We also examined the interplay between ensemble size and the number of denoising steps (**Fig. S10**). For the radius of gyration, the error with respect to our Mpipi-GG derived mean is flattened above 25 steps and 400 conformations. We, therefore, chose these values as default values for STARLING ensemble generation in our software implementation. However, we emphasize that larger ensembles may be necessary depending on the specific order parameter of interest.

Since R_e_ and R_g_ are highly correlated with sequence length, we next assessed the accuracy of our model on a set of length-matched sequences ^37^. This approach allowed us to determine whether the model had effectively captured the influence of sequence chemistry on the global dimensions of IDRs, or whether the model had simply learned to relate sequence length to global dimensions. Using all 100-, 200-, and 300-residue sequences with distinct sequence chemistries from the held-out test set, we found excellent agreement across a range of R_g_s (**Fig. 2C**, RMSE = [0.64 Å, 0.79, 1.06] and R^2^ =[0.99, 0.996, 0.996] for [100, 200, 300] residue lengths) and R_e_ (**Fig. 2D**, RMSEs = [2.7 Å, 3.5 Å, 4.4 Å], R^2^ = 0.98, 0.99, 0.99] for [100, 200, 300] residue lengths). These results gave us confidence that STARLING had learned *bona fide* sequence-to-ensemble rules instead of simply learning polymer scaling theory.

In addition to ensemble-averaged observables, full structural ensembles enable us to calculate distributions of any observable of interest (**Fig. 2E**). We set out to assess how well STARLING-derived global dimension distributions match distributions from simulations. We used the Hellinger distance as a similarity measure to compare distributions. The Hellinger distance ranges from 0 to 1, where zero indicates that the distributions are identical, and one signifies that they are entirely disjoint (**Fig. 2F**). The STARLING-derived R_g_ distributions show excellent agreement with distributions from simulations (**Fig. 2G**). This overlap is quantified by a low Hellinger distance, confirming the high degree of similarity between the two distributions.

Finally, rather than focusing on distributions from global descriptors (e.g. R_g_, R_e_), we took the more stringent approach to assess distributions of all possible inter-residue distances between simulations and STARLING ensembles. Here, we evaluated the Hellinger distance between every pair of inter-residue distances between Mpipi-GG and STARLING ensembles, again finding excellent agreement (**Fig. 2H, F**). In summary, all evidence supports that STARLING can directly predict meaningful conformational ensembles of disordered proteins from sequence.

Having established that STARLING can recapitulate simulated ensembles, we next investigated agreement with experimental data (**Fig. 3A**). We took a previously curated set of 133 sequences for which high-quality Small-angle X-ray Scattering (SAXS) data has been assembled, and average R_g_ values obtained. We found excellent agreement between average R_g_ values from STARLING with SAXS-derived R_g_, obtaining values that are comparable to the state-of-the-art from simulations (RMSE = **4.72**, R^2^ = **0.9**) (**Fig. 3B**, **Fig. S11**).

**Figure 3.**
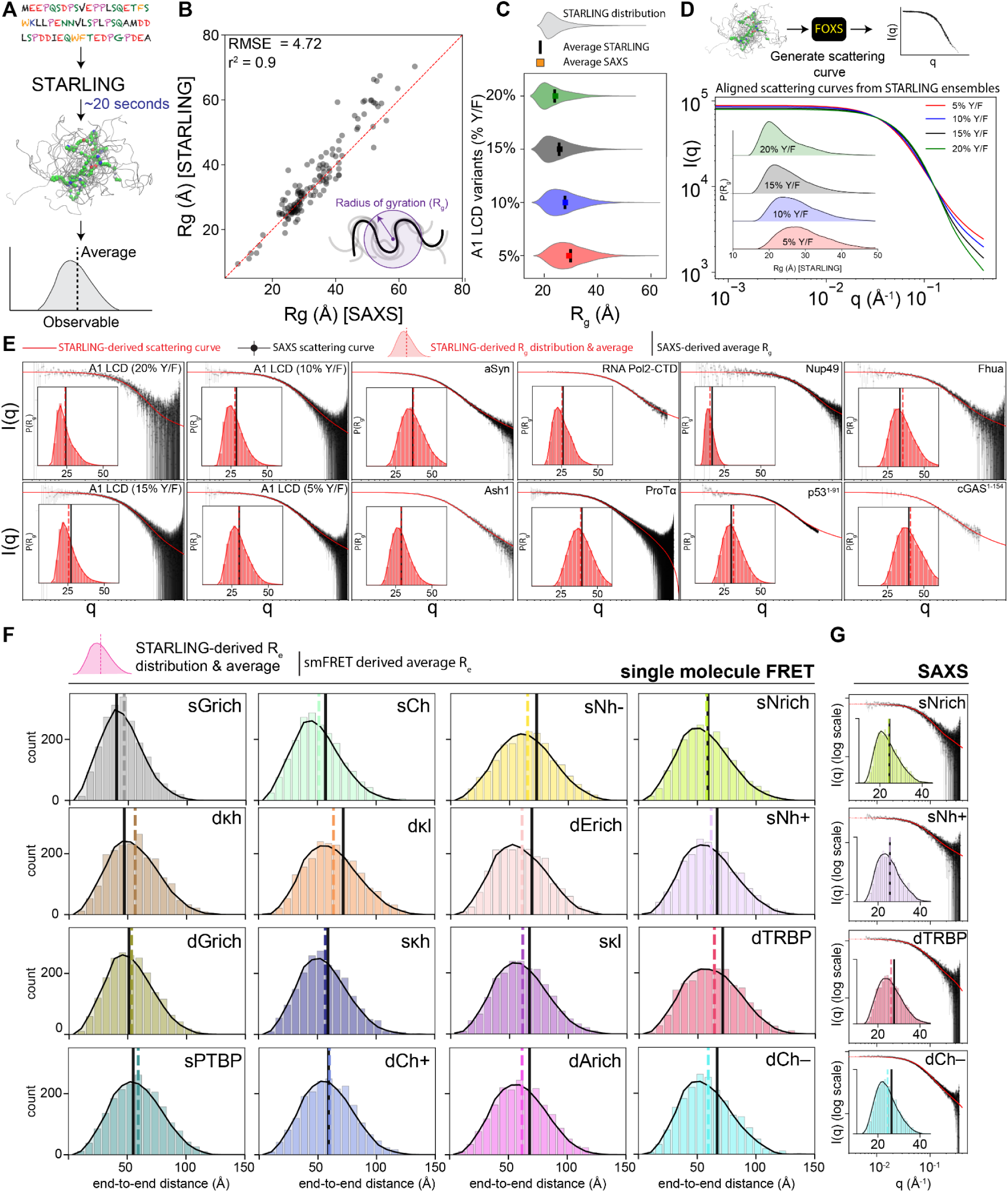
**(A)** STARLING enables rapid generation of ensembles from sequence. **(B)** STARLING shows state-of-the-art accuracy in terms of predicting the average R_g_ from sequence. 133 IDR sequences comparing SAXS-derived and STARLING-derived average R_g_ values (see also **Fig. S11**). **(C)** STARLING accurately captures the effect of small numbers of mutations, here illustrating how altering the number of aromatic residues changes the global dimensions of a low-complexity domain (LCD) taken from the RNA binding protein hnRNPA1. **(D)** Full scattering curves can be back-calculated from STARLING ensembles using FOXS, illustrating how changes in conformational behavior alter both global dimensions (small angles, low q value) and the shape of the ensemble (intermediate and high q values). **(E)** Comparison of twelve different full small-angle X-ray scattering (SAXS) profiles with scattering profiles calculated from STARLING ensembles. Insets show STARLING Rg distributions with average R_g_ values from scattering data shown as a solid black line. See also **Fig. S13-14**. **(F)** STARLING-derived R_e_ distance distributions with experimental values are shown as black lines for sixteen different length-matched disordered protein sequences with different sequence chemistries. See also **Fig. S15. (G)** SAXS scattering curves for unlabeled proteins for a subset of those in pane F.

Next, we evaluated the sensitivity of our model to capture previously established trends between IDR sequence chemistry and ensemble dimensions. To do this, we predicted ensembles for all length-appropriate IDRs in DisProt^38^, generating ensembles for 3,203 IDRs (**Fig. S12**). This revealed established sequence-ensemble relationships reported elsewhere (e.g., more compact IDRs are enriched in aromatic residues, whereas more expanded IDRs are enriched in proline and glutamic acid)^9,20,39,40^. To assess if STARLING can capture how small changes in sequence affect the ensemble, we generated ensembles for four variants of the hnRNPA1 low-complexity domain (A1-LCD) that vary the number of aromatic residues. These ensembles confirmed a clear relationship between aromatic content and dimensions, in quantitative agreement with the SAXS-derived experimental data (**Fig. 3C**)^9^.

While we initially investigated SAXS data by comparing it against R_g_ values (**Fig. 3B, 3C**), SAXS experiments provide scattering data that reports on polymeric properties of IDRs across a range of lengths (down to 5-10 Å resolution). Given STARLING generates full ensembles, we can use existing tools for back-calculating scattering curves to reconstruct synthetic scattering data from our ensembles, enabling direct comparison between raw experimental data and STARLING-derived conformations. Using FOXS, we first showed that the subtle changes seen for the A1-LCD variants are clearly observable in the resulting scattering profiles (**Fig. 3D**)^9,41^. We, therefore, took twelve unrelated IDRs where high-resolution scattering data were available and compared scattering profiles derived from STARLING ensembles to experimental data, finding excellent agreement across a wide range of sequence chemistries and sequence lengths (**Fig. 3E**). We further examined an additional forty scattering profiles, finding the vast majority are in excellent agreement with STARLING-derived scattering profiles (**Fig. S13-14**). Taken together, our data suggests that our STARLING ensembles faithfully reproduce coarse-grained descriptions of IDR ensembles across a range of length scales and sequence chemistries.

Finally, we sought to compare against an alternative experimental modality. Recent elegant work illustrated how a series of sixteen length-matched sequences show a range of end-to-end distances depending on the underlying sequence chemistry, providing the most comprehensive experimental and computational investigation into sequence-ensemble relationships to date^17^. We compared predicted end-to-end distances (and, where available, SAXS-derived scattering data) for those sixteen sequences, again finding good agreement between STARLING ensembles and experimental data and outperforming other approaches (**Fig. 3F, Fig. S15, Fig SXX**). Of note, our average error here across the sixteen sequences (RMSE 6 Å) is comparable to the differences obtained between different experimental groups in a recent smFRET benchmarking study^42^. Overall, our work suggests that STARLING can generate full coarse-grain ensembles of IDRs that accurately capture experimental data.

Finally, we applied STARLING to investigate a set of distinct IDRs in various contexts. We aim to illustrate the types of investigation STARLING enables rapidly and efficiently. The master transcriptional regulator cMyc (439 residues) underlies cell growth, metabolism, and proliferation and is frequently dysregulated in many cancers^43^. cMyc consists of a large N-terminal IDR (cMyc^1–361^), a C-terminal HLH DNA binding domain (cMyc^368–406^), and a leucine zipper (cMyc^413–439^). The large size of its IDR coupled with its chemical composition has challenged biophysical characterization (**Fig. 4A**). Using STARLING, we generated conformational ensembles of cMyc^1–361^ to investigate sequence-encoded conformational biases in the ensemble (**Fig. 4B, Fig. S16**). In particular, we were interested to determine whether conformational ensembles offered insight into regions within cMyc known to bind other partners (so-called Myc boxes). Curiously, the STARLING-derived ensemble suggests that cMyc^1–361^ can be divided into two halves (IDR1 and IDR2). IDR1 (cMyc^1–200^) is more compact and engages in intramolecular interactions and some long-range interactions. IDR2 (cMyc^201–361^) is more expanded with locally compact regions. Finally, conformationally distinct subregions align with established Myc-box boundaries, indicating local and long-range conformational properties can align with established functional annotations. Overall, these results predict cMyc^1–361^ contains strong sequence-encoded biases that may be correlated with (or even underlie^44^) its function.

**Figure 4.**
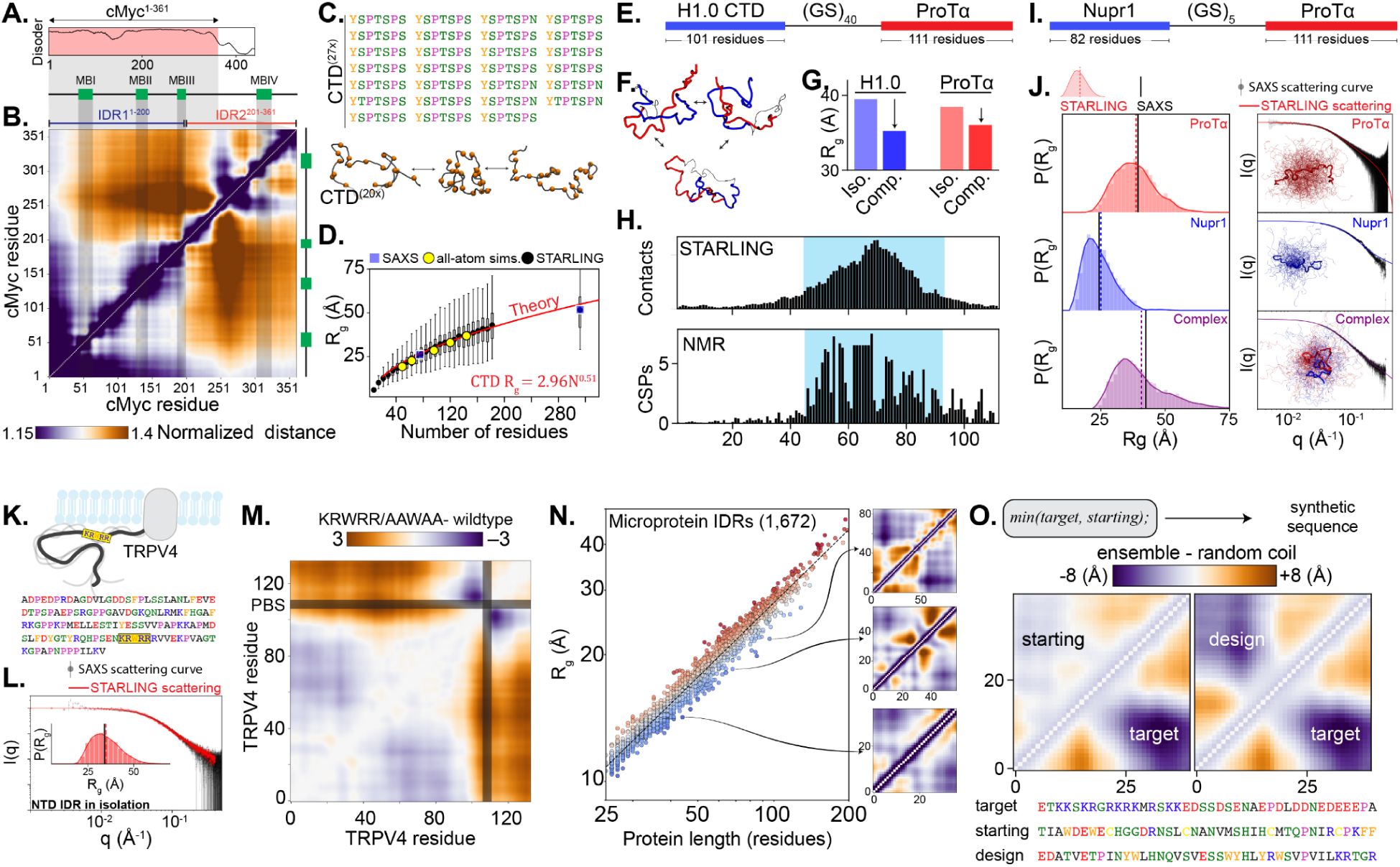
**(A)** The master transcriptional regulated cMyc is largely disordered. **(B)** STARLING provides a full description of the cMyc^1–361^ conformational ensemble, identifying distinct subregions and residues that drive attractive and repulsive intra-molecular interactions. Inter-residue distances here are normalized by a limiting polymer model for a polypeptide as a Gaussian chain; darker colors are closer together, and lighter colors are further away. **(C)** The RNA polymerase II CTD consists of a series of heptad repeats. **(D)** STARLING ensembles show excellent agreement with SAXS and all-atom simulations, enabling polymer theory to describe CTD dimensions with a simple analytical expression. **(E)** Histone H1.0 and ProTα form a high-affinity complex together. We generated ensembles for a construct in which the H1.0 CTD and ProTα are tethered by an inert (GS)_40_ linker. **(F)** The resulting ensemble shows robust H1.0:ProTα interaction. **(G)** H1.0CTD and ProTα contract in the complex vs. in isolation. **(H)** Comparison of per-residue ProTα H1.0-CTD contacts and NMR-derived chemical shift perturbations (CSPs) show excellent agreement. **(I)** The proteins Nupr1 and ProTα have also been shown to form a complex, and we took a similar approach with a shorter (GS)_5_ linker to enable reasonable characterization comparison to SAXS data. **(J)** STARLING-derived ensembles show excellent agreement with Nupr1, ProTα, and the Nupr1:ProTα complex. **(K)** The TRPV4 channel has a large N-terminal IDR with a basic PIP_2_ binding site (PBS). **(L)** STARLING-derived ensembles show good agreement with small-angle X-ray scattering data. **(M)** The PBS engages in long-range interactions with a distal acidic region, and mutation of positive residues reduces these long-range interactions, in good agreement with extant experimental work. **(N)** Many microproteins are small disordered proteins; therefore, STARLING enables systematic prediction of microprotein ensembles. Note that the y-axis is transformed via log10 **(O)**. STARLING enables the rational design of sequences to match a desired distance map. Here, a random starting sequence is dissimilar to the target sequence, and after rounds of optimization, a new, entirely distinct sequence that recapitulates the ensemble behavior of the target sequence is generated. Distances here are shown as absolute inter-residue distance with a random coil model subtracted.

Although STARLING enables the investigation of hitherto unexplored IDRs, it can also provide new insight into existing systems. The C-terminal domain (CTD) of the DNA-directed RNA polymerase II subunit RPB1 consists of a series of heptad repeats with the consensus sequence YSPTSPS **(Fig. 4C)**^45^. This region stretches for 52 repeats in humans, and the number of repeats varies across organisms. Given the repetitive nature of the CTD, we reasoned principles from polymer physics would be well-suited to quantitatively describe the global dimensions as a function of CTD length. To this end, we generated STARLING ensembles across 26 different CTD lengths to generate a systematic series. We also performed all-atom Monte Carlo simulations across five different lengths^46,47^. The all-atom and STARLING derived ensembles show excellent agreement with one another as well as extant SAXS data **(Fig. 4D)**. Fitting the radius of gyration vs. number of amino acids to a polymer scaling law of the form R_g_ = B_0_N^ν^ (where R_g_ is the radius of gyration, B_0_ is a prefactor term that captures a combination of chain stiffness and monomer excluded volume, and ν is the apparent Flory scaling exponent) revealed well-fit parameters of B_0_ = 2.96 Å and ν = 0.51. The convergence of STARLING ensembles, all-atom simulations, SAXS, and theory all suggest that – at least with respect to *in vitro* conditions – the global dimensions (including end-to-end distance, **Fig. S17**) can be reasonably well-captured using a simple analytical description, enabling the 3D dimensions of CTDs from across the kingdoms of life to be estimated directly.

We next wondered if we could use STARLING to characterize disordered bimolecular complexes. In the early days of AlphaFold2, biomolecular interactions were investigated by tethering two folded domains using a flexible glycine-serine (GS) linker. We reasoned a similar approach might be possible with STARLING **(Fig. 4E)**. Tethering the C-terminal disordered region from the Histone H1.0 (H1.0^97–198^) to the histone chaperone Prothymosin α (ProTα) enabled the formation of an electrostatically-driven complex between the two proteins **(Fig. 4F)**, as reported previously^15^. Both proteins contract in the complex **(Fig. 4G)**, as observed and predicted by theory^48^. Moreover, the specific residues in ProTα identified by NMR-derived chemical shift perturbations (CSPs) match those residues from the STARLING complex ensemble that drive intermolecular interactions **(Fig. 4H)**. These results indicate that STARLING-derived binary complex ensembles can capture elements of IDR:IDR interaction ensembles.

While there is a limited number of biophysically-characterized disordered complexes, we also investigated the complex between ProTα and the fully disordered 82-residue protein Nupr1 (**Fig. 4I**). Prior work has characterized the proteins individually and in complex using SAXS^49^. We again find good agreement between STARLING ensembles and SAXS data for the individual proteins and – using a shorter GS linker (GS) – the disordered bi-molecular complex (**Fig. 4J**). Overall, our work suggests that STARLING has the potential for the prediction of fully disordered bi-molecular complexes, enabling the development of precise sequence-specific molecular hypotheses that can be tested experimentally.

Although our analyses thus far have primarily focused on ensemble differences between very different sequences, we wondered if STARLING could interpret relatively small amino acid changes. The N-terminal IDR from the Transient Receptor Potential Vanilloid 4 (TRPV4) ion channel has been thoroughly characterized through a battery of biophysical techniques to reveal a regulatory long-range interaction between a basic PIP_2_ binding site and an acidic cluster (**Fig. 4K**)^50^. STARLING-derived ensembles of this region show good agreement with SAXS data (**Fig. 4L)**. Moreover, comparing the difference in intramolecular distances for the wildtype vs. a KRWRR-to-AAWAA mutant that was shown experimentally to reduce long-range interactions, we observe a commensurate reduction in long-range interactions in STARLING **(Fig. 4M)**. These results illustrate that STARLING enables the investigation of even relatively modest changes in sequence to examine sequence-ensemble-function relationships.

We were also interested in investigating STARLING’s application to larger-scale ensemble prediction. Recent work has characterized a large set of microproteins – non-canonical ORFs typically under 100 amino acids in size – using a combination of mass spectrometry and Ribo-Seq-based approaches^51^. While these microproteins are highly disordered, they possess distinct amino acid sequence biases from ‘canonical’ ORF-derived IDRs **(Fig. S18**). From 1785 microproteins in these data, we identified 1672 disordered regions and predicted ensembles for all of them. While many short IDRs are relatively expanded, we noticed many sequences with extensive intramolecular contacts driven primarily between aromatic and arginine residues (**Fig 4N**). These results suggest that even short-disordered proteins can possess sequence-encoded conformational biases.

Finally, given the speed with which STARLING enables ensemble prediction, we wondered if we could use STARLING to generate sequences that match the conformational ensemble of a target sequence. To this end, we developed an ‘ensemble dissimilarity metric’ based on STARLING-generated distance maps and used it as part of an objective function for the design of IDRs with desired ensemble properties (see *Methods*). As a proof-of-concept, we started with a randomly generated amino acid sequence and used STARLING to optimize the sequence towards matching the conformational ensemble of a short region in the C-terminal IDR of CTCF (Uniprot ID: P49711, residues 590-629). We generated a new sequence that matches the long and short-range behaviors of CTCF^590–629^ **(Fig. 4O, Fig. S19)**. Alongside similar methods^36,52,53^, the ability to rationally design IDRs with desired conformational properties opens the door to test the importance of conformational biases in disordered protein ensembles.

Overall, these vignettes demonstrate the versatile range of applications STARLING enables. Notably, the speed and ease with which STARLING generates ensembles means these investigations can be accomplished in hours or even minutes.

## DISCUSSION & CONCLUSION

In this work, we present STARLING, a multiscale generative model capable of producing coarse-grained IDR ensembles directly from protein sequence in seconds. While there are many caveats and limitations (discussed below), the primary advantage we see with STARLING is that it drastically reduces the barrier to obtaining conformational predictions on sequence-dependent IDR behavior. While IDRs are often ignored or visualized in terms of AlphaFold “orange spaghetti,” STARLING makes it straightforward for anyone to obtain realistic coarse-grained ensembles. Rather than “the end” of an investigation into IDR sequence-ensemble behavior, we view STARLING as a beginning, making it straightforward to develop hypotheses as to how an IDR’s sequence may determine its conformational ensemble and/or how it may influence interactions with other IDRs.

STARLING does not produce dynamical trajectories (i.e. there is no time component), but instead enables the rapidly generating large ensembles. While this may be seen as a weakness, we see this as a strength in that the slow reconfiguration times of large chains often hampers simulations from obtaining a sufficient number of truly independent conformations. In contrast, STARLING ensembles for long IDRs are generated just as quickly as small ones, and because conformers are generated via independent stochastic denoising processes, there is no configurational correlation between consecutive conformers to consider.

A key feature of STARLING is the ability to generate full conformational ensembles instead of predicting just ensemble average values. While predicting average conformational behaviors (e.g., average R_g_ or average R_e_) can be informative^20,21,54^, it does not enable a ‘structural’ investigation into the determinants of a given behavior. With a conformational ensemble, the specific sets of residues responsible for an observed behavior can be directly identified (**Fig. S20-21**). Moreover, substantial effort has been placed into developing methods to back-calculate experimental observables from simulation trajectories to interpret experimental data^55^. STARLING ensembles are amenable to these analyses, allowing comparison with a range of experimental results.

Our decision to focus on coarse-grained ensembles over all-atom ensembles was driven by several factors. Firstly, obtaining sufficient training data (especially to capture emergent long-range behaviors for larger IDRs) at all-atom resolution for larger IDRs would be prohibitively difficult with current approaches. While we could, in principle, back-map from coarse-grained to all-atom conformations, whether such back-mapping accurately reproduces local conformational biases at all-atom resolution is unclear, given the limitations of an isotropic amino acid bead. Moreover, it is not apparent whether sufficient experimental data exists to validate (or refute) such an approach.

While STARLING generates coarse-grained conformational ensembles within seconds, it does have a number of limitations which we believe are important to acknowledge and highlight. Firstly, the ensembles generated by STARLING lack any information about secondary structure. Since STARLING models are trained on data from one-bead-per-residue coarse-grained force fields, we are blind to putative secondary structural elements in the disordered protein sequence. Secondly, STARLING operates on single-chain, fully disordered protein sequences; however, disordered regions are generally found either adjacent to and/or between folded domains^1^. While STARLING is powerful at recapitulating the solution behavior of strictly disordered protein sequences, there is emerging biophysical evidence that folded domains (or membranes) can influence the biophysical ensembles of adjacent disordered protein sequences. Although a limitation of the raw predictive power of STARLING, we view this as a “feature” rather than a “bug” as deviations from conformational ensembles obtained in isolation could indicate biophysically meaningful interactions (and potentially function). Moreover, with enough compute, careful design, and consideration for sequence library design and structural coverage, future models could address this via massive large-scale simulations of structurally diverse folded domains tethered to physicochemically disparate disordered sequences.

STARLING models are trained on data generated from simulations performed under “standard conditions”; however, an increasingly appreciated role for intrinsically disordered proteins is their involvement in environmental sensing in response to changes in the local chemical environment^56^. The increased solvent accessibility of disordered proteins and unique physicochemical sequence biases poises IDRs to be particularly responsive to changes in this local chemical environment (e.g., salt, pH, metabolites, nucleic acids, other proteins, etc.)^57–61^. Understanding how disordered proteins respond to these changes in the solution environment is thus an important area of future research. Since STARLING was not trained on these data types, its reliability in this field cannot be determined. Moreover, since the field lacks good experimental data to benchmark these models, it is unclear how useful a deep-learning-based surrogate model would be for these use cases, which starkly contrasts the typical solution behavior we rigorously compare against in this work. Nevertheless, similar models could be trained with more advanced simulation models as the field progresses.

Another limitation of STARLING is its inability to guarantee that generated conformations are Boltzmann-distributed. While scoring functions learned by DDPMs have been shown to approximate the underlying force field, these models do not sample directly from the canonical ensemble^62^. Nevertheless, it’s recently been shown that the scoring function learned by a DDPM is a relatively strong approximation of the underlying force fields^62^. Indeed, in some cases, this scoring function can actually be employed as a force field to directly sample conformers from the approximate force field^62^.

Several other approaches for generating IDR ensembles from sequences have been released over the last year. The first example of applying DDPMs to IDR ensembles came from foundational work by Taneja and Lasker, who offered compelling proof-of-concept that a two-stage approach, akin to AlphaFold2, could be used to learn sequence-to-ensemble mapping for chemically heterogeneous sequences^63^. idpSAM is a latent-space DDPM that enables C-alpha predictions trained on 3,259 all-atom Monte Carlo simulations^64^. However, idpSAM, in principle, offers predictive power limited to sequences that are up to 60 amino acids in length, although in practice, we found poor agreement with experimental data even within this range (**Fig. S22**). IDPFold is another approach for ensemble generation that combines protein language models with experimental and molecular dynamics data^65^. However, our evaluation of IDPFold-derived ensembles against experimental data yielded relatively poor results, with ensemble prediction times taking substantially longer and scaling poorly in terms of runtime and memory requirements with sequence length (**Fig. S23-S24**). Finally, recent work has focussed on developing machine learning models for predicting Hamiltonians (as opposed to the raw training data)^66^, while others have taken generative models trained on folded proteins and shown these models can reproduce unfolded behavior consistent with disordered proteins^67^. Overall, however, extant tools lack the accuracy, performance, and ease of install/use that STARLING provides.

A promising direction for future research is the integration of generative models—conditioned on various force field parameters—into a multi-fidelity Bayesian optimization framework to improve force field design^68,69^. Currently, developing force fields involves an iterative process that combines experimental data and simulations to identify a parameter set that closely matches the experimental observables. However, this approach is hindered by the need for sequential tuning of force field parameters and costly numerical simulations to evaluate their accuracy. To accelerate this process, researchers may be able to leverage cheaper surrogate models, such as the latent generative model proposed here (low-fidelity), to rapidly scan the force field parameter space and identify promising regions of hyperparameter space. These candidate regions can then be validated using more intensive high-fidelity simulations, allowing for a more efficient exploration of the parameter landscape. The ultimate goal of this approach is to use a computationally efficient methodology, combining conditional generative models with targeted high-fidelity simulations, to discover optimal simulation parameters that might have been inaccessible through traditional methods. More work will be needed to address how force field parameters can be encoded to make generative modeling effective, transferable, and physically realistic ^62,70,71^.

Despite recent advancements in structural biology methodologies (e.g., cryoEM and cryoET), key structural information is often missing from essential proteins involved in cellular signaling and information processing. This is because these structure-centric techniques struggle to capture intrinsically disordered regions, which are highly flexible and, therefore, incompatible with traditional structure-centric approaches. The absence of this information limits our understanding of biophysical mechanisms in these essential pathways. STARLING presents an exciting opportunity to serve as putative restraints in larger integrative structural models to address this challenge^72^.

In conclusion, we hope STARLING will be useful to the disordered protein community and beyond. In contrast to many deep learning tools, we have made a substantial effort to ensure STARLING is easy to use and install (e.g., weights are managed automatically, installation is as easy as pip install, and prediction can be done using a single command-line command). The ease with which STARLING facilitates biophysical investigation into IDR conformational behavior is substantial. Coarse-grained ensembles can be predicted on the fly on a whim, in meetings, or during seminars, while large-scale analysis can be performed on commodity hardware in hours. While the validity of predictions should always be considered, and we do not see STARLING as a replacement for any existing technologies, we nevertheless see STARLING filling a currently unmet need in enabling biophysical investigation of IDR behavior to the broader community.

## Author Contributions

**Conceptualization:** BN, JML, ASH; **Methodology**: BN, JML; **Software**, **Validation**, **Formal analysis**, **Investigation**: BN, JML, RJE, ASH; **Resources & Funding Acquisition**: JML, ASH; **Data Curation**: BN, JML, ASH; **Writing & Visualization**: BN, JML, RJE, ASH;

## Acknowledgements

We thank Jackie Pelham, Ramiz Somjee, and Bede Portz for their comments on the manuscript, and we thank Ute Hellmich for giving such an exceptional talk that inspired our investigation of the TRPV4 IDR. We especially thank our many colleagues for sharing and depositing raw experimental data (notably SAXS data on SASDB), which has greatly facilitated comparison with experimental data. We also thank Jungsan Sohn, Scott Showalter, and Maria Grazia Ortore for sharing raw SAXS scattering data. We thank Birthe Kragelund and Freia Stryhn Buus for sharing chemical shift data for the ProTa:H1.0 complex. We thank Erik Martin and Ben Schuler for sharing SAXS data of the linker sequences.

## Version comment

We note that this version of the manuscript lacks supplementary tables with all of the sequences used in this study. All these data will shortly be available at https://github.com/holehouse-lab/supportingdata/tree/master/2025/starling_2025.

## METHODS

### Sequence design and data splitting

We needed to construct a physicochemically diverse library of sequences to train our models for downstream simulations/model training. The sequence library presented in this work encompasses an expanded set of the constructs originally presented in Lotthammer *et al*.^20^. In addition to those sequences, we have also added 4,284 tri-block copolymer sequences (2994 in training, 654 in test, and 636 in validation). These tri-block sequences consist of three blocks: N and C terminal blocks of particular chemistry (consisting of a repetitive low complexity sequence) separated by a comparatively inert block. The N and C terminal blocks may be the same but generally differ. For example, one potential sequence would be positive charges, an inert GS linker, and negative charges.

After designing the sequence library and prior to model training, we clustered the sequences using MMseqs2^73^ protocol (**`mmseqs easy-cluster input.fasta output_directory tmp --min-seq-id 0.7 -c 0.8 --cov-mode 1`**). Next, we stratified the cluster centers by length and assigned all sequences within each cluster into either train (49,423 sequences, 70%), validation (10,703 sequences, 15%), or test sets (10,437 sequences, 15%) (**Fig. S1**).

### Molecular Dynamics Simulations and Analysis

Unless otherwise noted, we followed the same simulation protocol described previously^20^. Briefly, this process included performing Mpipi-GG simulations with the LAMMPS simulation engine^74^. We built initial configurations of all disordered proteins according to a random walk. For all simulations, we performed up to 1000 iterations of energy minimization until the force tolerance was below 1 x 10^-8^ (kcal/mol)/Å. Simulations were performed in the NVT ensemble at 150 mM NaCl. All simulations employed a weakly-coupled Langevin thermostat that was adjusted every 100 ps to maintain a target temperature of 300 K. The integration timestep was 20 femtoseconds for all simulations. Periodic boundary conditions were used in all simulations with a box length of 500 Å. All simulations were performed for 10 µs with trajectory coordinates saved every 2 ns.

All of our simulations were analyzed with SOURSOP and MDTraj^75,76^. For all simulations analyses, we chose to subset the data for (1) computational tractability and (2) to reduce the correlation between conformers by striding our analysis to only compute inter-residue distance maps every 30 frames, equating to one conformation for every 60 ns or 3 million integration steps. Collectively, we have a total of 9,621,465 inter-residue distance maps.

### Generative Model Training

We coupled two deep learning methodologies, a variational autoencoder^77^ (VAE) and a denoising-diffusion probabilistic model^30,78^ (DDPM), for efficient sequence-to-ensemble predictions. We constructed training, validation, and test datasets using coarse-grained simulations performed with the Mpipi-GG force field. Our dataset consists of sequences between 10 and 384 amino acids in length, and due to this variability in length, we padded all our distance maps to a consistent size of 384 by 384.

We trained a variational autoencoder to learn a compressed latent representation of distance maps. We chose to train a VAE to represent our distance maps in a 24 by 24 latent space, which amounts to a spatial compression ratio of 256. These design decisions enabled us to reduce the computational resources required for model training and inference while retaining strong performance. Following standard practices in VAE model training, our training objective consists of a reconstruction loss and a Kullback-Leibler (KL) divergence term. The reconstruction loss is modeled as negative log-likelihood under a Gaussian distribution, and KL loss regularizes the latent space by penalizing deviations from a unit Gaussian distribution (**Fig. S3**). We down-weighted the KLD loss by a factor of 1e^-4^, similar to previously reported approaches^31^, to permit our model to balance accurate reconstruction of inter-residue distance maps with a regularized and structured latent space.

Next, we developed a latent diffusion model by training a discrete time Denoising-Diffusion Probabilistic Model (DDPM) within the latent space of a variational autoencoder (VAE). The DDPM is a generative model that learns the data distribution by gradually denoising samples starting from isotropic Gaussian noise over a series of timesteps. To achieve this, we first corrupted the data through a noising process (i.e., the forward process) where Gaussian noise is progressively added according to a noise schedule.

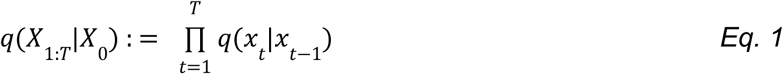

where T ranges from t=1 to t=1000 and 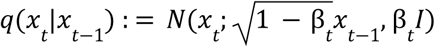, where beta is defined by the variance scheduler described below.

To ensure robust learning of the noise added at each time step – randomly sampled from discrete values between t=1 (original data) and t=1000 (fully noised data) – we employed a variance schedule that controls the magnitude of the added noise throughout the diffusion process. The schedule begins by adding a small amount of noise (low variance) where the original data is “information dense” and gradually increases. In our case, we used a cosine scheduler to select variances between 0.0001 and 0.999 with an offset factor of 0.008. This offset has a minor effect and is primarily motivated by addressing pixel binning issues in image models, where data is constrained to a [0,255] dynamic range. However, this rationale does not directly translate to our spatial dataset, which is unnormalized and has different structural properties. While the impact of this offset in our case is expected to be minimal, we acknowledge its suitability for our data is unverified. We did not conduct ablations to assess its effects, and alternative approaches tailored to our dataset could be worth exploring in future work. A neural network was then trained to reverse this process conditioned upon the current timestep and the noisy data at that time step.

To parameterize a generative model, we train a neural network to reverse the time-dependent addition of Gaussian noise described in the forward process above. The reverse process is defined by the following relation:

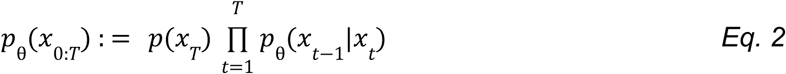

where *p*_θ_ (*x*_*t*−1_ |*x*_*t*_) is a Gaussian distribution, and *p*_θ_ (*x*_*t*−1_|*x*_*t*_) = *N*(*x*_*t*−1_; µ_θ_ (*x*_*t*_, *t*), β_*t*_ *I*) and µ_θ_ (*x*_*t*_, *t*) is computed from the predicted noise ɛ (*x*, *t*) according to the following equation:

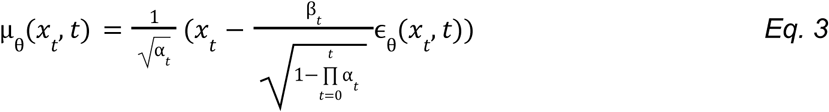

The parameter β_*t*_ is fixed according to the time-dependent variance noise scheduler described below and α_*t*_ : = 1 − β_*t*_. The next denoised state, *x*_*t*−1_, is given by *x*_*t*−1_ = µ_θ_ (*x*_*t*_, *t*) + σ ɛ where

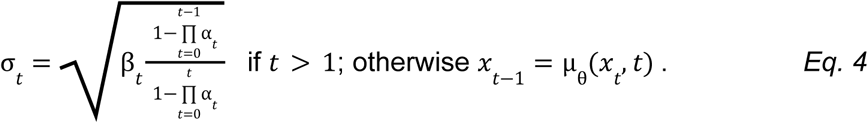

We utilize a U-Net architecture with both residual convolutions and transformer blocks. The model processes batches of 2D isotropic Gaussian noise with a single feature channel and spatial dimensions concordant with the VAE latent space (**Fig. S5**). The input undergoes three steps of spatial compression and feature extraction, increasing the feature count by 512x and reducing the spatial dimensions by 64x at the base of the U-Net. This is followed by a series of upsampling blocks that gradually restore the spatial dimensions and reverse the feature extraction process, ultimately predicting the noise content within the original input. Prior to each downsampling and upsampling convolutional step, we augment our residual blocks with a transformer. Each transformer includes both self-attention, where each pixel in the latent space attends to all others, and cross-attention, which captures the relationship between the learned sequence embeddings and the latent representation of distance maps. The sequence embeddings are a (512, ℓ)-dimensional vector where ℓ is the 384 residues. If a given sequence is less than 384 residues, we pad the input sequence to a length of 384. The 512-dimensional vector comprises both learned amino acid token embeddings and incorporated positional encodings via the addition of sinusoidal embeddings to those token embeddings. Our sinusoidal embeddings use the commonly used theta parameter of 10,000. Taken together, incorporating these cross-attention layers enables our model to conditionally generate sequence-specific distance maps.

Finally, the diffusion model is trained with a mean squared error loss function between the added and predicted noise according to the following equation:

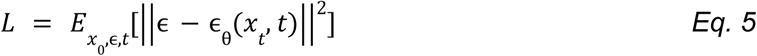

During training, distance maps are encoded using a pre-trained VAE, with the output rescaled to achieve unit variance. Noise is then added to the latent representation, and our transformer-augmented U-Net predicts the amount of noise introduced.

### Model inference

Putting it all together, during inference, 2D isotropic Gaussian noise alongside sequence information is iteratively input through the trained U-Net until all the noise is removed. The final output of the U-Net is then fed into the VAE decoder to project back to a ℓ x ℓ matrix of inter-residue distances consistent with the input sequence. Importantly, each instance of the 2D isotropic Gaussian noise is sampled independently, allowing us to generate decorrelated IDR conformations. The iterative process of removing noise can be time-consuming and typically requires thousands of iterations through the neural network. Custom samplers aim to reduce the number of passes through the neural network while maintaining high-quality samples by redefining the forward and reverse process of the DDPMs. Following advances in the image generation literature, we have implemented a denoising-diffusion implicit model^79^ (DDIM) to speed up the generation of samples from our model. Since DDIMs share the same training objective as DDPMs, no additional training is required, and we can use DDIM samplers to improve the inference speed of any trained DDPM.

### Projecting distance maps to structural ensembles

To project from 2D inter-residue distance maps back to 3D coarse-grained structures with the same set of inter-residue distance distributions, we use a custom PyTorch implementation optimized for our use case of the Scaling by MAjorizing a COmplicated Function (SMACOF) algorithm - we note that we did not choose this approach for its apt name (although we endorse it, as it follows the spirit of our software package names), but rather for its improved rate of convergence on stress optimization. We note that in our preliminary tests, our implementation offers a ∼70x improvement over the default SciPy implementation.

If PyTorch-compatible hardware (MPS or CUDA) acceleration is not available, we provide a fallback to the scipy implementation of the Multidimensional Scaling (MDS) algorithm available in sklearn (*sklearn.manifold.MDS*). Briefly, this algorithm works by projecting our 2D inter-residue distance matrices - where each entry d_ij_ in the input matrix represents the spatial distance between residues i and j - into 3D cartesian coordinate feature space. When performing this projection, we use these inter-residue distances as the precomputed dissimilarity measure. This projection aims to derive a set of 3D coordinates that best capture the spatial relationships between residues, as reflected in the input distance map.

Mathematically, this projection optimizes a stress function that quantifies the discrepancy between the pairwise distances in the 2D matrix and those in the projected 3D space. The stress function is defined as:

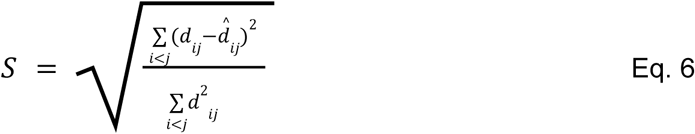

Where d_*ij*_ is the distance between residues *i* and *j* in the original 2D distance matrix. *d̂_ij_* is the corresponding Euclidean distance between residues *i* and *j* computed from the projected 3D space.

### Intrinsically Disordered Protein Ensemble Design

To enable IDR sequence design with target biophysical ensemble properties, we use the sequence optimization functionality from the Python package GOOSE (https://github.com/idptools/goose)^36^. Briefly, this sequence design approach iteratively modifies a starting sequence using random amino acid substitutions to minimize the mean squared error between a property or set of properties computed (or predicted) from a sequence. For the sequences designed in this manuscript, we leverage STARLING and this optimization approach to design novel sequences by computing an ensemble dissimilarity score. To compute this ensemble dissimilarity score, we use STARLING to generate distance maps for the target ensemble. Next, we take the sequence at the current iteration of the optimization process and compute the ensemble of distance maps for this sequence. Then, we compute the mean and standard deviations of the distances for each residue pair in the distance maps for both sequences. To account for variations in distance due to inter-residue proximity, these values are normalized by dividing by the distance from the diagonal. The sum of the absolute difference between the target sequence and the second sequence is then calculated for the normalized mean and standard deviation maps. The function then calculates the absolute difference between the target sequence and the current sequence for both the normalized mean and standard deviation maps. Finally, it combines these differences into a single scalar value that represents the dissimilarity between the two sequences.

The EnsembleDesign property comes with many tunable parameters that can be initialized and adaptively set during sequence optimization for improved performance. This primarily entails a progressive increase in the number of denoising steps and the number of conformations used as the initial sequence progresses closer and closer to the target ensemble. We also provide a patience functionality that controls how long optimization can continue without improving before increasing the number of steps or conformers.

For sequence optimization, we started with a randomly generated sequence. For our EnsembleDesign property, we set the conformations to always be 200 and the minimum and maximum number of steps to be 5 and 25, respectively. We also set the number of rounds of optimization without sequence improvement before increasing the number of steps to be 500 iterations. At each iteration, the starting sequence was randomly changed by substituting a random number of a randomly chosen amino acid for some other amino acid in the starting sequence. Next, the difference between the new sequence and the target sequence was quantified as described above. This evolutionary algorithm continues until convergence or the stopping criteria have been met. We opted to replace cysteine residues in our final designs with serines, although this constraint could easily be encoded into the design algorithm by preventing cysteine sampling.

### Distribution overlap with Hellinger distance

To compute the Hellinger distance between two distributions, we use the implementation provided in the SOURSOP **`sssampling`** module, as described previously^80^. Briefly, this involves generating a density-normalized histogram for the two distributions of interest with the same bin values, and then measuring the overlap between those histogram densities (P and Q) as described below:

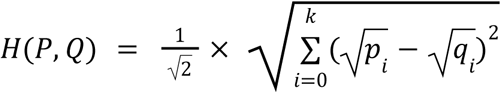

To ensure parity, all distributions are generated using identical numbers of datapoints, although in principle, this is not a prerequisite for this approach to work.

### Comparison with SAXS profiles

SAXS profiles are back-calculated using FOXS with default parameters and using 400 conformers (water_layer_c2 = 2.0, excluded_volume=1.0, max_q=0.4)^41^. The profiles were aligned by selecting a short region of experimental and computational profiles and identifying a scalar that would transform the computationally-derived I(q) values to match the experimentally measured I(q) values in that region. This allows us to superimpose the two curves on top of one another using common I(q) values (where are reported in arbitrary units) but with the measured q values (reported in Å^-1^). Many examples of scattering data were obtained from SASBDB^81^. For the 133 SAXS sequences used in the comparison in Fig. 3, these were taken from prior work, filtering for those sequences which were under 384 amino acids in length^20^. For the 52 SAXS sequences for which scattering curves were compared, we sought out high-quality scattering data from reported systems in the literature.

### Large-scale ensemble predictions and bioinformatics

For DisProt prediction, we used DisProt obtained in the fall of 2024 (**`DisProt_release_2024_06_with_ambiguous_evidences.fasta`**) obtained from DisProt^38^. We filtered for IDRs between 25 and 380 amino acids in length. This resulted in 3,204 distinct IDR sequences.

For microprotein analysis, microprotein IDRs were predicted using metapredict V3. The reference human proteome was obtained from UniProt (**`UP000005640_AND_revi_2025_02_07`**) and all IDRs predicted using metapredict V3^2,82^. This analysis identified 33,473 IDRs in the human proteome. Compositional enrichment was calculated over all residues in distinct datasets.

### All-atom simulations

All-atom simulations for CTD repeats of length 49, 63, 96, 120, and 144 amino acids were performed using the CAMPARI simulation engine and ABSINTH implicit solvent model using the ion parameters of Mao *et al.* as described previously ^9,83–85^. Simulations were performed at 380 K to mitigate challenges in sampling (as described previously for aromatic-rich low-complexity sequences^9^), using 10 mM NaCl, with five independent replicas performed for 6 x 10^7^ Monte Carlo steps each, with the first 1 x 10^7^ steps discarded for equilibration. Conformers were saved every 2 x 10^4^ steps, meaning each independent simulation generated 2500 conformations. Overall, for each CTD length, we obtained atomistic ensembles of 12,500 conformers. Atomistic ensembles were analyzed using SOURSOP^76^.

### Null model normalization

To enhance the visualization of inter-residue distances, we normalized inter-residue distances using the Analytical Flory Random Coil (AFRC) model^86^. This model represents the expected dimensions for a sequence-specific version of a polypeptide if the protein behaved as a true Gaussian (phantom) chain; that is, no excluded volume and no inter-residue attraction or repulsion. This becomes useful as a reference point to normalize the effects of sequence separation when comparing ensembles.

### STARLING software implementation and installation

The STARLING software package is built using Python 3.11. STARLING makes heavy use of SciPy^87^, NumPy^88^, and PyTorch^89^. Documentation is built using Sphinx and hosted on ReadTheDocs. Testing is enabled by PyTest. Structure manipulation is facilitated by MDTraj ^75^.

#### Model weights

Initial weights are tagged as v1.0.0 and available at https://github.com/idptools/starling/releases/tag/v1.0.0. STARLING handles weights automatically on the backend, such that the first time a new install of STARLING is run it will automatically download the model weights for the VAE and DDPM. This happens once and requires an internet connection. After the weights have been downloaded once, STARLING defaults to using the cached versions found in **`∼/.cache/torch/hub/`**. From the user perspective, this provides a seamless install experience.

#### Installation

To install, we recommend using conda to set up an isolated environment, although we note STARLING has no “esoteric” dependencies. To install, simply run;

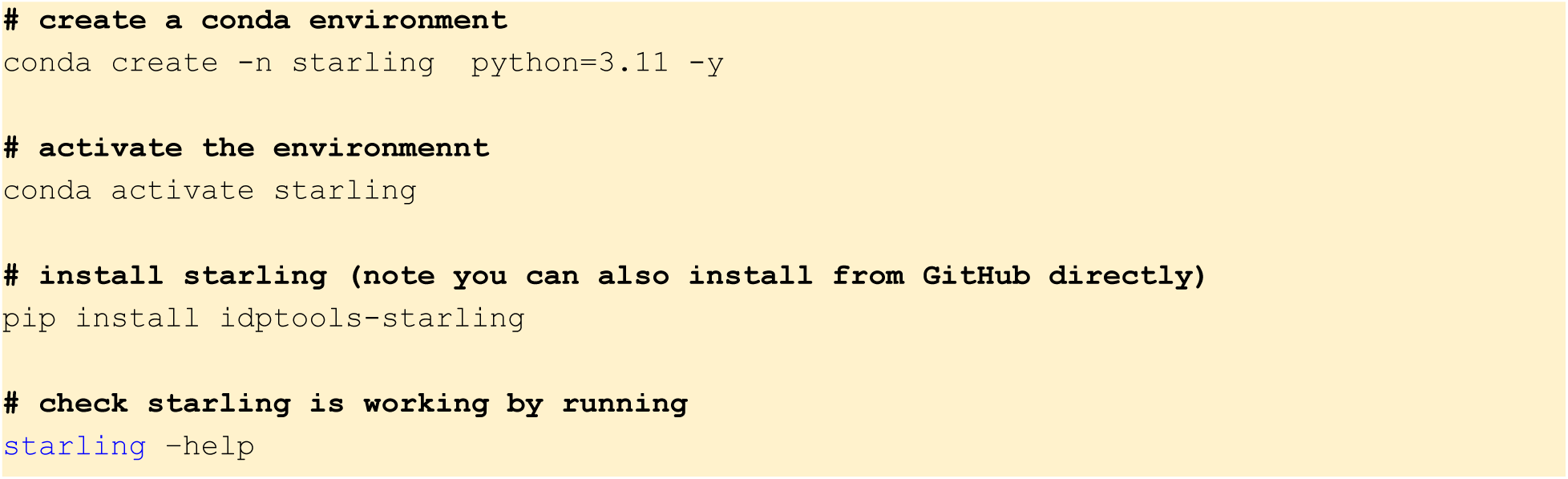

Note that the first time any starling command is run it may take 30-90 seconds to initialize various things in your new Python environment and download the STARLING weights in the background. This is all a one-time thing.

#### Command-line tools

To generate an ensemble with starling, simply run

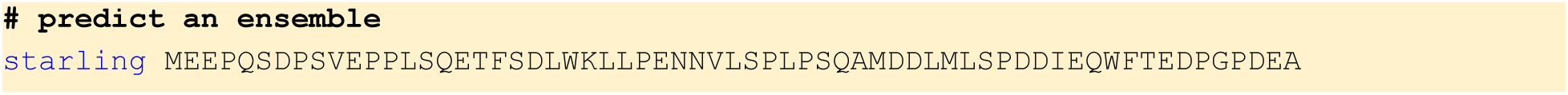

To define the number of conformers, use the `-c` (or `--conformations`) flag. By default, the name of the output file generated if a single sequence is passed will be `sequence_1.starling`. To set a custom name for the output file, use the `--outname` argument. The **`starling`** command can take a single sequence or a FASTA file with multiple sequences. In this situation, the names of the output files will be defined by their FASTA headers. For a complete set of documentation run

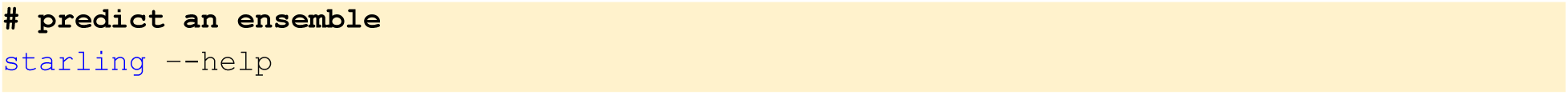

By default, STARLING generates `.starling` files as output. These files contain all the data and metadata associated with the ensemble prediction. To visualize the ensemble as a set of molecular structures, you can convert `.starling` files into PDB trajectories (or XTC/PDB pairs) using our conversion tools:

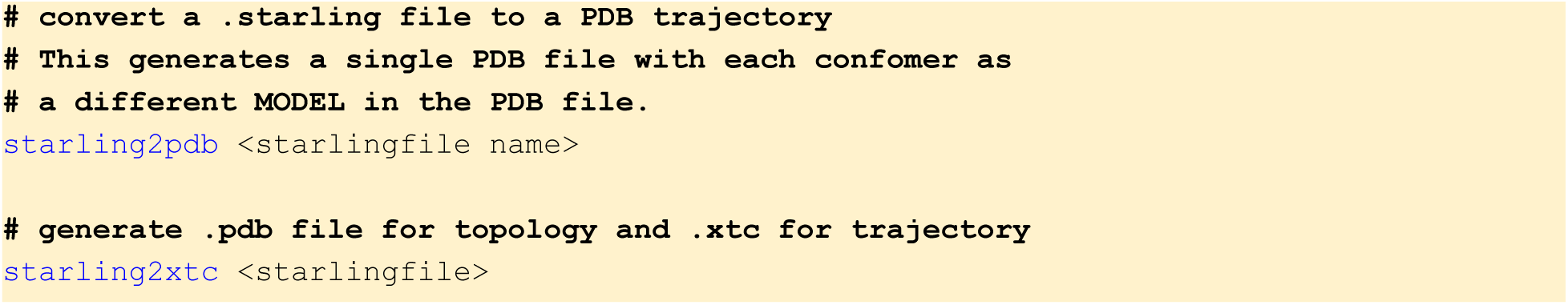

#### Python library

When analyzing STARLING ensembles we highly recommend using the STARLING Python library instead of converting ensembles to structural ensembles and then analyzing them in this context. This is because converting to structural ensembles requires structure reconstructing, which can introduce (minor) errors. In contrast, analysis of the predicted distance maps mitigates those errors.

Working with STARLING ensembles in Python is straightforward. STARLING has just two user-facing functions: **`generate()`** and **`load_ensemble()`**.

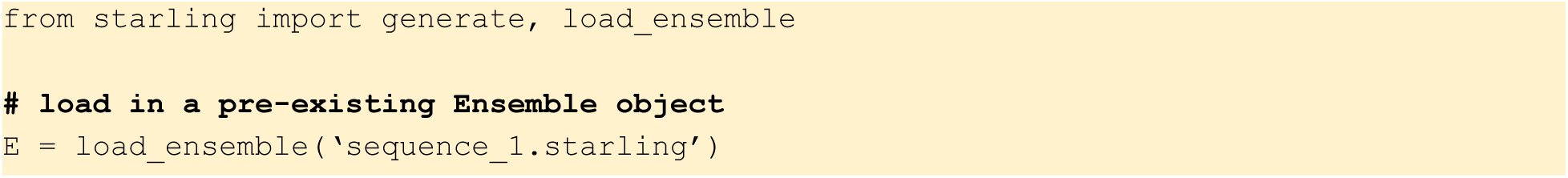

The Ensemble object has a range of functionality enabling the investigation of ensemble properties. New Ensemble objects can also be generated using the **`generate()`** function.

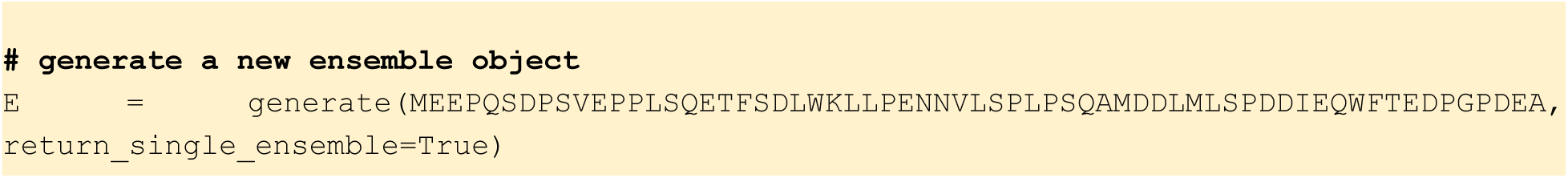

The documentation and development for this is ongoing; for feature requests or errors, please raise an issue at https://github.com/idptools/starling/issues.

### Compute Hardware

All model prototyping was performed on an in-house deep learning workstation comprising 4 NVIDIA A4500s. All production training runs were performed on NVIDIA GH200 GPUs provided via generous support courtesy of the Texas Advanced Computing Center and the Frontera Computational Sciences Fellowship.

### Code distribution

STARLING is fully open source and provided here: https://github.com/idptools/starling/

STARLING’s documentation is available here:

https://idptools-starling.readthedocs.io/en/latest/

STARLING is provided as a Colab notebook here: https://github.com/idptools/idpcolab/blob/main/STARLING/STARLING_demo.ipynb

The code and data associated with the figures in this manuscript will (shortly) be available here (we promise!):

https://github.com/holehouse-lab/supportingdata/tree/master/2025/starling_2025

