## Supplementary Information for "Accurate predictions of conformational ensembles of disordered proteins with STARLING"

VERSION 1.1 (2025-02-24)

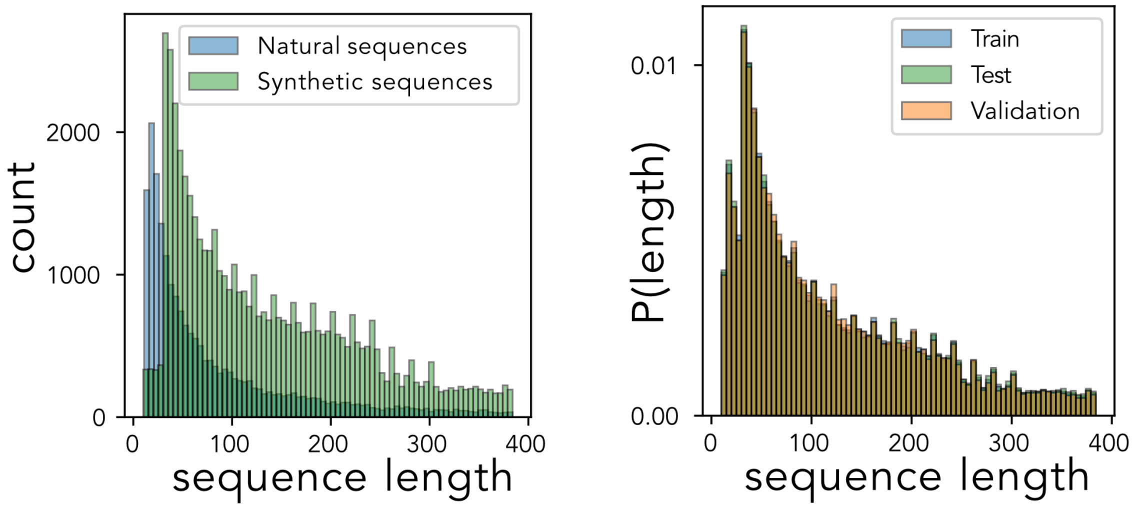


**Figure S1:** Our dataset of 70,563 consists of both natural sequences (N=20,349) and synthetic sequences designed by varying sequence features known to affect IDR ensembles (N=50,214) (left). The sequences were clustered and split into train (N=49,423), validation (N=10,703), and test (N=10,437) sets while maintaining the distribution of sequence lengths across the splits consistent (right).

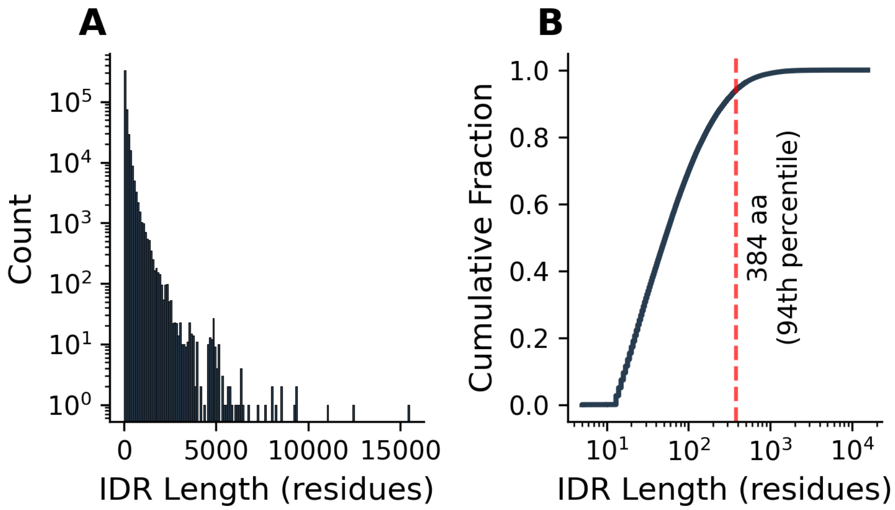


**Figure S2:** Disordered protein length distributions across a collection of model organism proteomes. **A.** Histogram of the protein sequence length distribution. **B.** A cumulative density function of the length distribution presented in A. 94% of the IDRs across common model organisms fall between 0 and 384 residues (the length cutoff of STARLING).

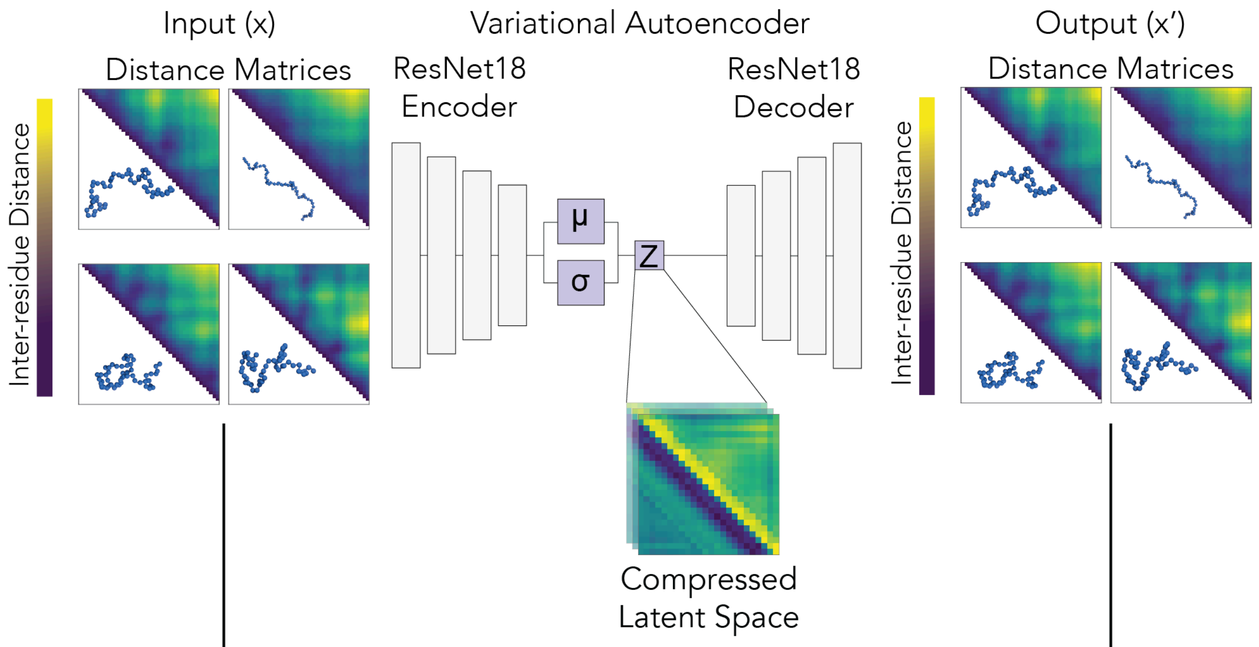


$$\mathcal{L} = \mathbb{E}_{q_\phi(z|x)}\left[\log p_\theta(x \mid z)\right] - D_{\mathrm{KL}}\left(q_\phi(z \mid x) \| p(z)\right)$$

where $p_\theta(x \mid z) = \mathcal{N}\left(\mu_\theta(z), \sigma^2 I\right)$ and $x' \sim p_\theta(x \mid z) = \mathcal{N}\left(\mu_\theta(z), \sigma^2 I\right)$

**Fig. S3:** Schematic representation of the variational autoencoder (VAE). The input to our variational autoencoder is a collection of 2D inter-residue distance maps that undergo feature extraction and spatial compression through a ResNet18 encoder, which learns the μ and σ parameters of a normal distribution that parameterizes the latent features. The VAE is trained with an evidence-based lower bound (ELBO) loss comprising a reconstruction term and a Kullback–Leibler (KL) divergence term.

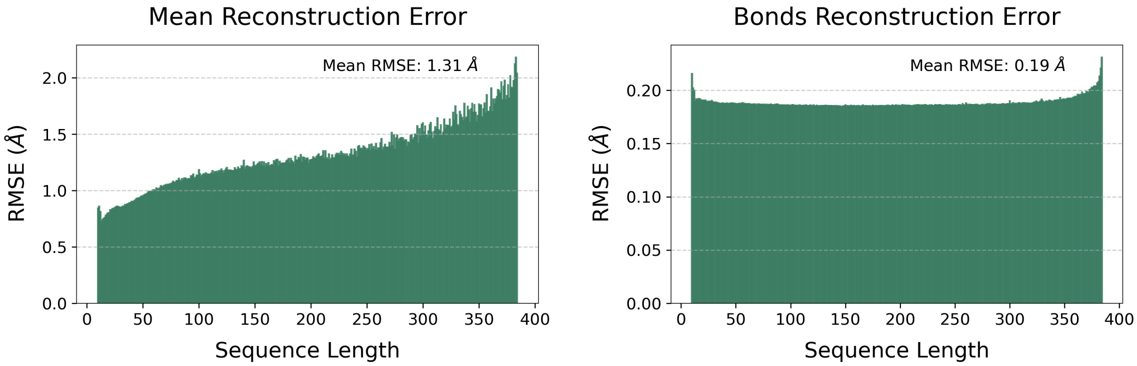


**Figure S4:** The variational autoencoder (VAE) is evaluated on a held-out test set of distance maps (N = 1,425,849). The VAE encoder spatially compresses these distance maps and is then reconstructed by the VAE decoder from their compressed representations. The root mean squared error (RMSE) is computed between the input and output distance maps. The trained VAE demonstrates a low mean RMSE, calculated over the upper triangle of each distance map (left). Accurate bond reconstruction is essential for generating physically realistic conformations, and our model achieves a very low mean RMSE (right).

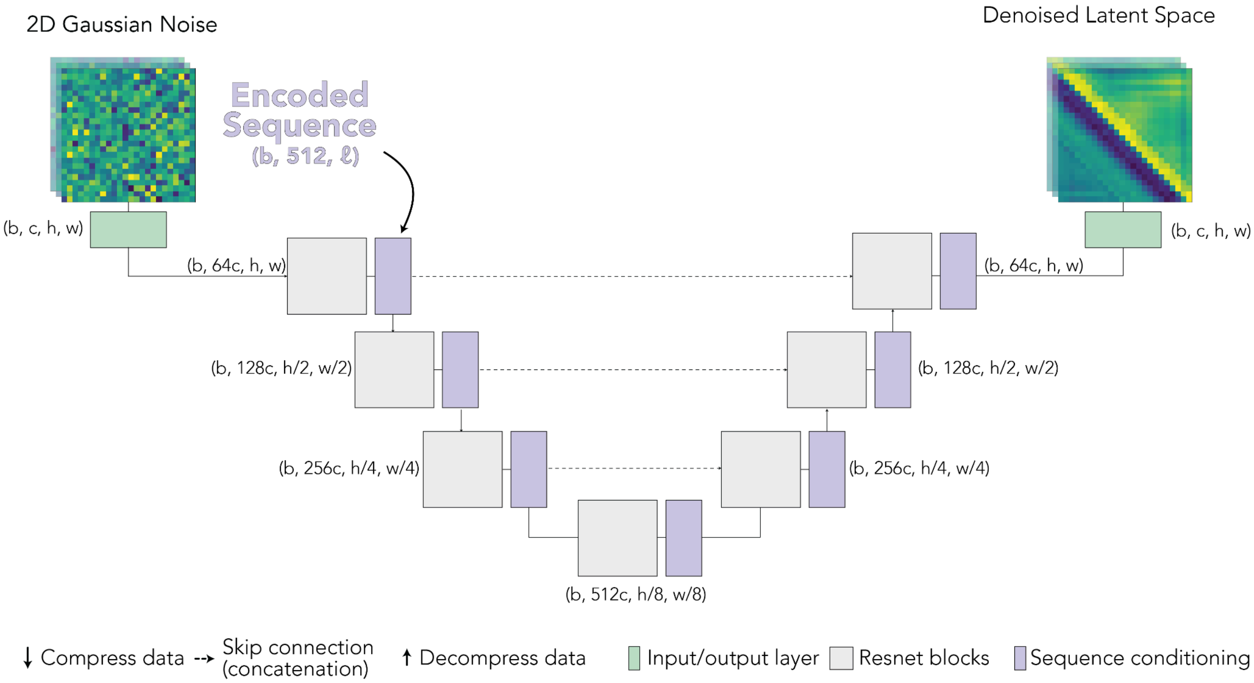


**Fig. S5:** Extended schematic representation of the Unet backbone of the STARLING discrete time denoising-diffusion probabilistic model (DDPM). The DDPM starts by taking a batch size (b) of 2D Gaussian noise with one channel (c) of features and spatial dimensions concordant with the VAE latent space (h,w). This data then undergoes three steps of spatial compression and feature extraction until we reach the base of the “U”, where we have 512x the features and 64x spatial compression (8x*8x). Following downsampling is a series of upsampling blocks that reverse the feature extraction and spatial compression, ultimately arriving at the original batch, channels, height, width dimensional vector (b,c,h,w) where the batch is the number of conformers, the channels are the noise estimate at each latent feature, the height and width are the compressed spatial dimensions which can subsequently be fed to the VAE decoder to project back to a ℓ x ℓ matrix of inter-residue distances. Notably, at each step of the downsampling and upsampling, we inject sequence information into the model by cross-attending to sequence embedding where each amino acid is encoded as a 512-dimensional vector comprising both learned amino acid token embeddings and positional encodings via the addition of sinusoidal embeddings.

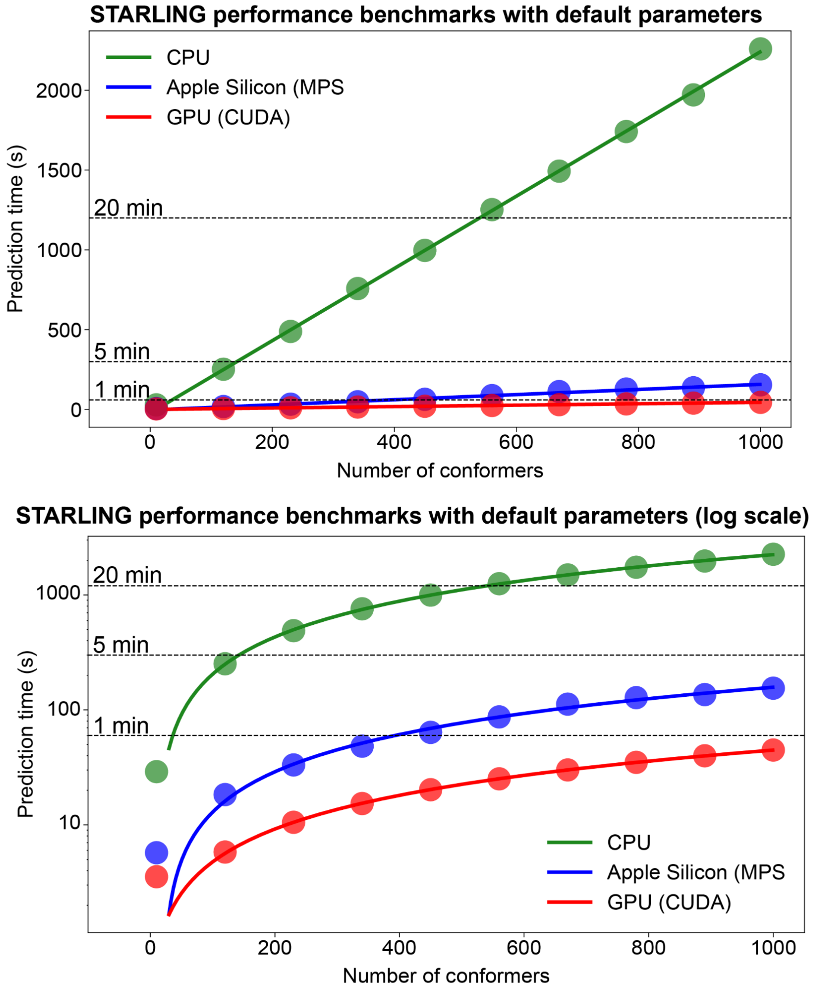


**Fig. S6.** STARLING benchmarking on different types of hardware. Runtime depends linearly on the number of conformers (although sequence length has no impact on runtime). The top and bottom panels show identical data in log vs. linear space. On GPUs and Apple Silicon (facilitated by CUDA and MPS, respectively), STARLING is highly performant, offering ensembles sufficient for detailed biophysical investigation in 10-15 seconds (GPU) or 30-60 seconds (MPS). Specific hardware tested here are Nvidia A4500 (GPU), Macbook Pro M3 Max CPU (MPS), and Intel(R) Xeon(R) Silver 4210R CPU @ 2.40GHz (CPU).

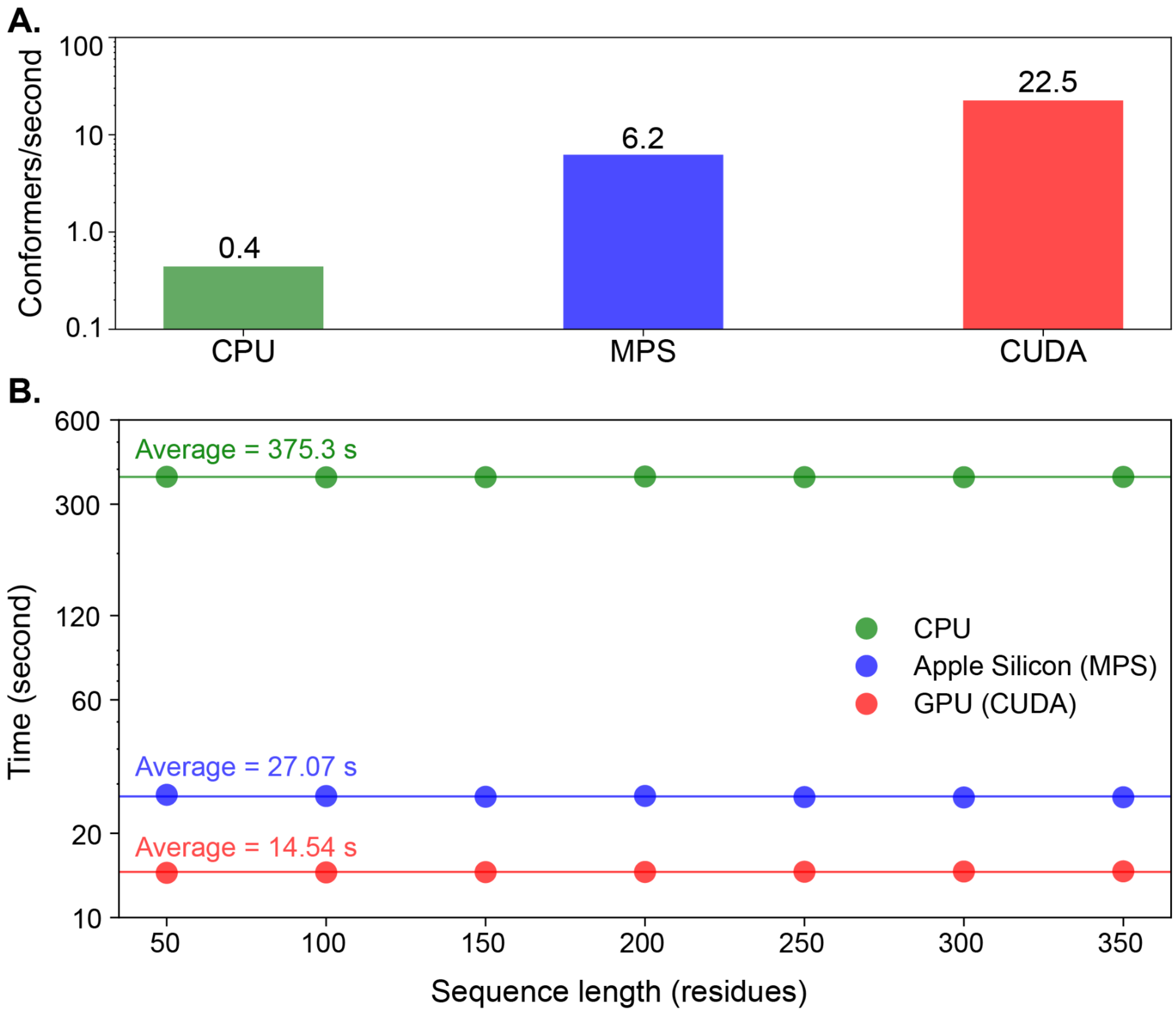


**Fig. S7. (A)** Number of conformers generated per second on different hardware types. Specific hardware tested here are Nvidia A4500 (GPU), Macbook Pro M3 Max CPU (MPS), and Intel(R) Xeon(R) Silver 4210R CPU @ 2.40GHz (CPU). **(B)** STARLING ensemble predictions show neither runtime nor memory scaling based on sequence length. Note here the GPU used was an NVIDIA RTX A4000, hence the slight drop in performance compared to panel (A).

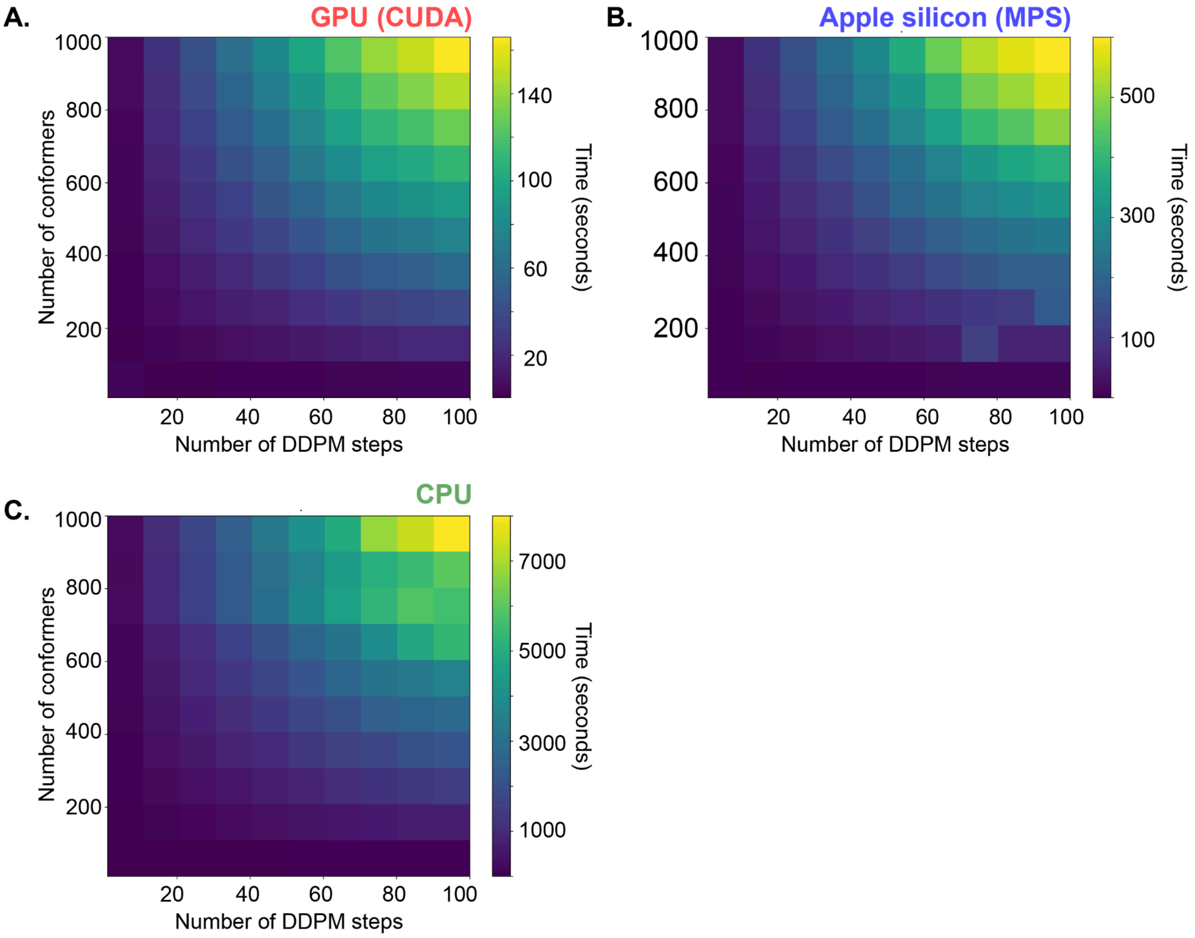


**Fig. S8.** Sweep of runtime vs. ensemble size and number of denoising steps on different hardware types. Specific hardware tested here are Nvidia A4500 (GPU), Macbook Pro M3 Max CPU (MPS), and Intel(R) Xeon(R) Silver 4210R CPU @ 2.40GHz (CPU).

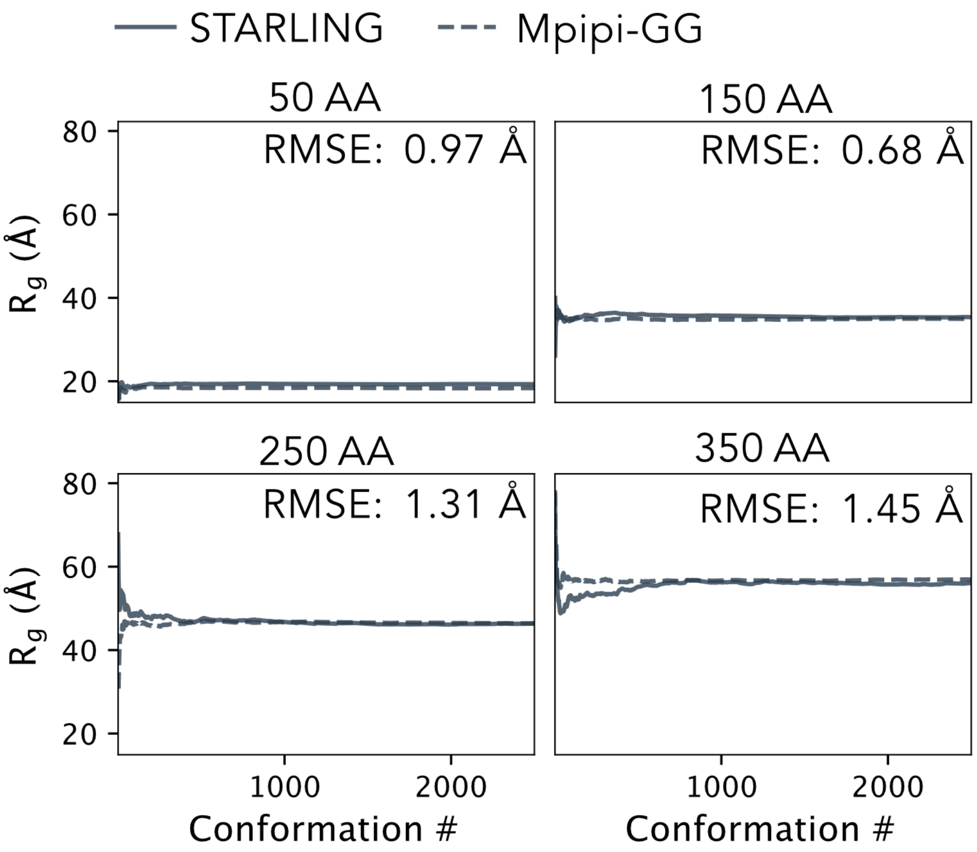


**Fig. S9.** Radius of gyration ($R_g$) convergence plot as a function of the size of the ensemble (number of conformations) for varied IDR lengths (50, 150, 250, and 350 residues in length). The dashed line is the ensemble-average $R_g$ value taken from long Mpipi-GG simulations. STARLING-derived $R_g$ values converge within the first 1000 conformers, exhibiting minimal error compared to Mpipi-GG ensembles. Based on this analysis, we selected 1000 STARLING conformers for $R_g$ prediction comparisons in all benchmarking used here.

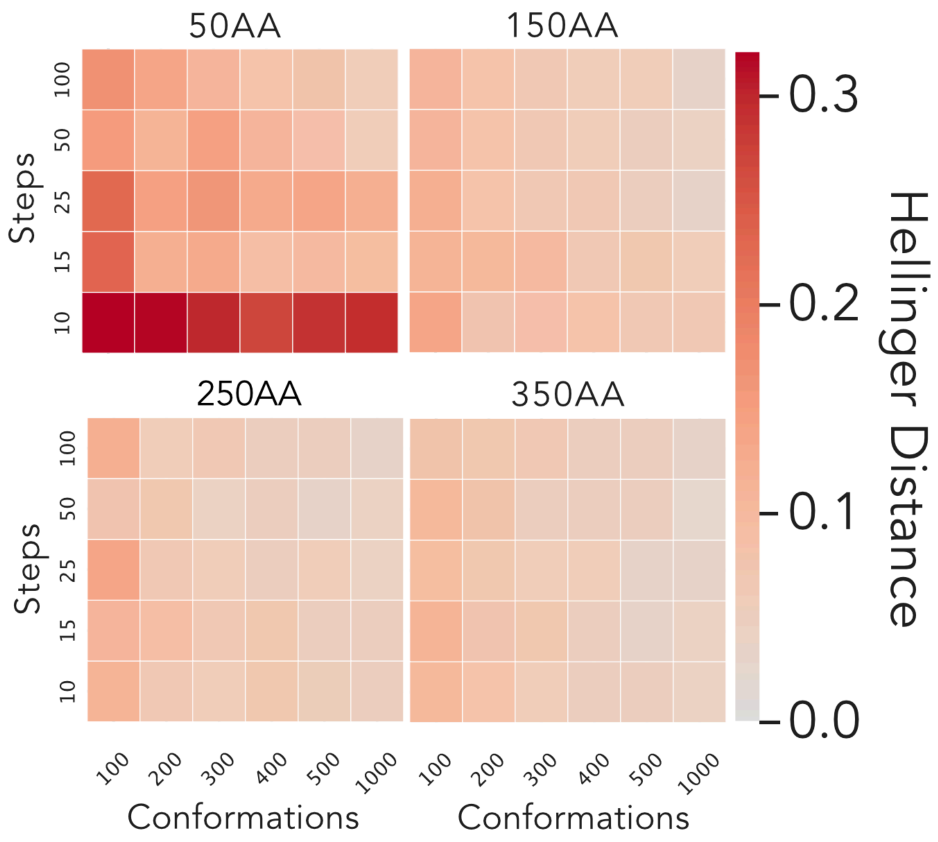


**Fig. S10.** Convergence plot depicting the relationship between Hellinger distance, the number of conformations sampled from STARLING, and the number of denoising steps for various IDR lengths (50, 150, 250, and 350 residues). The Hellinger distance quantifies the similarity between two distributions, where a value of zero indicates identical distributions and a value of one signifies no overlap (**Fig. 2F**). The Hellinger distance decreases as the number of conformations and denoising steps increase, with optimal STARLING performance observed at 1000 conformers and 25 denoising steps.

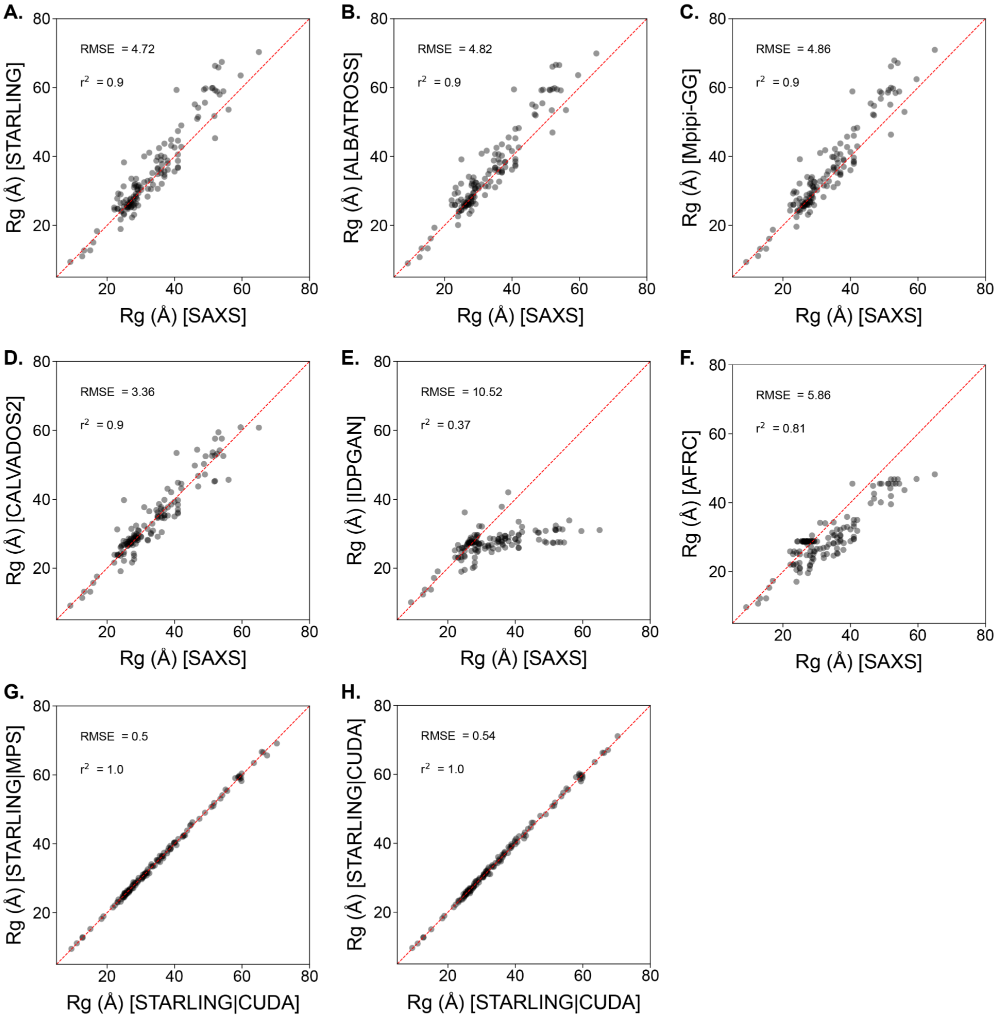


**Fig. S11.** Comparison of experimental $R_g$ values obtained from 133 different experimental datasets with various state-of-the-art tools for ensemble prediction: **(A)** STARLING (this work), **(B)** ALBATROSS, a deep-learning-based predictor that can only predict average ensemble values, **(C)** Mpipi-GG, coarse-grained molecular dynamics simulations, **(D)** CALVADOS2, coarse-grained molecular dynamics simulations, **(E)** idpGAN, a Generative Adversarial Network for predicting ensembles from the sequence, **(F)** The Analytical Random Flory Coil [AFRC], a limiting null model that considers IDPs as Gaussian-like chains with no interactions between residues. We include this as this is the true "lower bound" for what a model should achieve in terms of accuracy and RMSE. **(G)** We also compared $R_g$ predictions on GPU vs. MPS implementations for completeness. This revealed 1:1 agreement that matched the RMSE and correlation obtained when the same dataset was analyzed twice on the same hardware **(H)**.

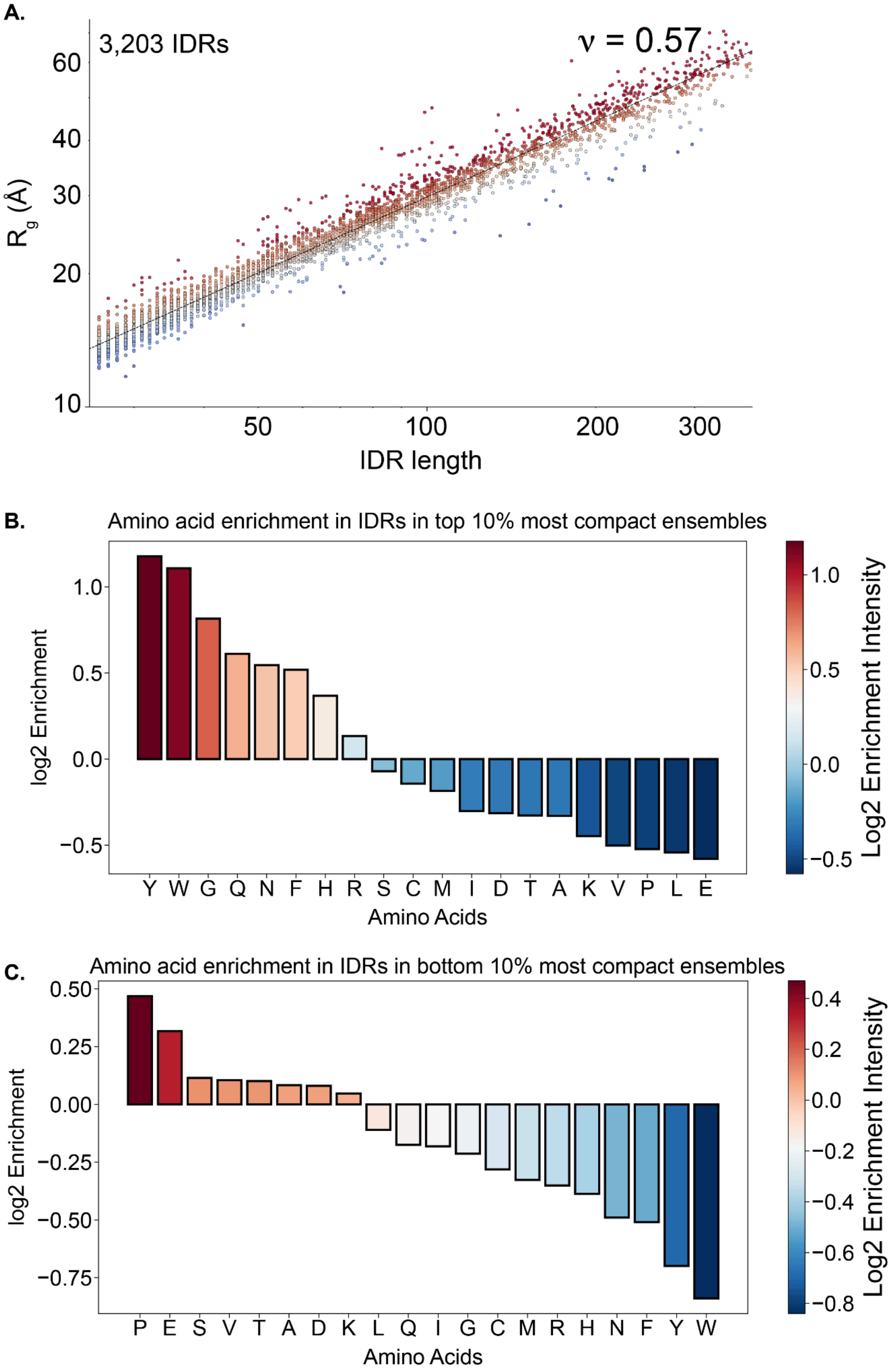


**Figure S12.** STARLING ensembles for DisProt. We predicted all ensembles in ~18 hours using a desktop computer with an NVIDIA A4500 GPU. **(A)** Fitting ensemble dimensions vs. length reveals an apparent scaling exponent of 0.57, indicating these IDRs are relatively expanded. However, we note an unavoidable acquisition bias in experimentally-characterized IDRs, meaning highly soluble (and hence more expanded, fewer intra/intermolecular interactions) are preferred for biophysical characterization. We also note that we underestimate the contributions of aliphatic hydrophobes in driving intramolecular interactions and any effect from local helicity or the long-range consequences of intramolecular helix interaction. As such, we strongly caution against interpreting the value of 0.57 to mean all IDRs are highly expanded. **(B)** Analysis of sequences in the most compact 10% of IDRs identified aromatic residues as being over-represented here, in line with prior work [1–3]. (C) An analogous analysis for sequences in the top 10% of expanded IDRs also reveals negatively charged residues and proline as being enriched in sequences that drive expansion, again, in agreement with prior work[1,4,5].

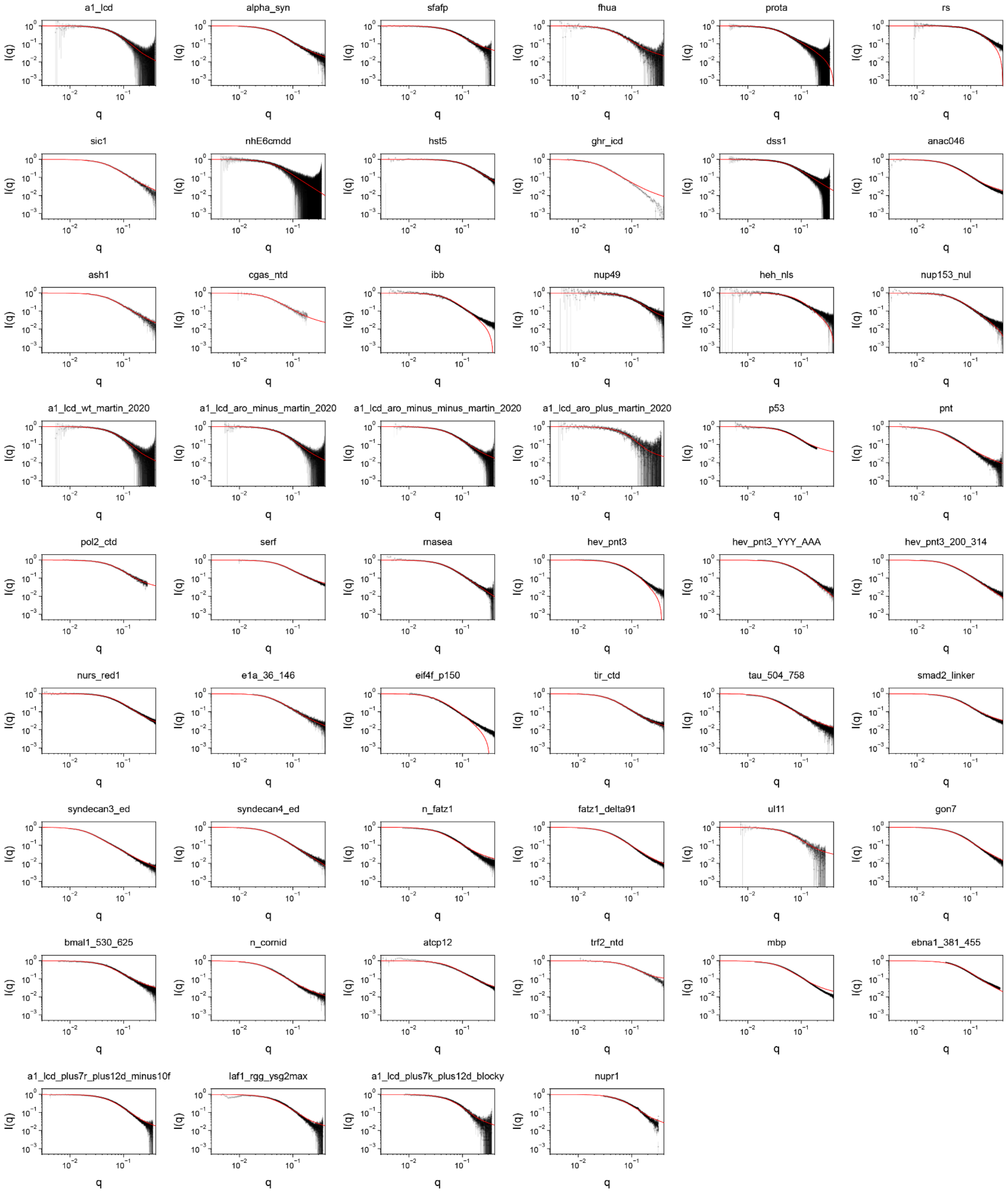


**Figure S13.** Comparison of STARLING-derived scattering data with experimental profiles from 52 sequences and datasets. Black is experimental and red in STARLING-derived scattering data. STARLING-derived curves were back-calculated using FOXS. We emphasize that this agreement is not a direct fit to experimental data but rather a prediction.

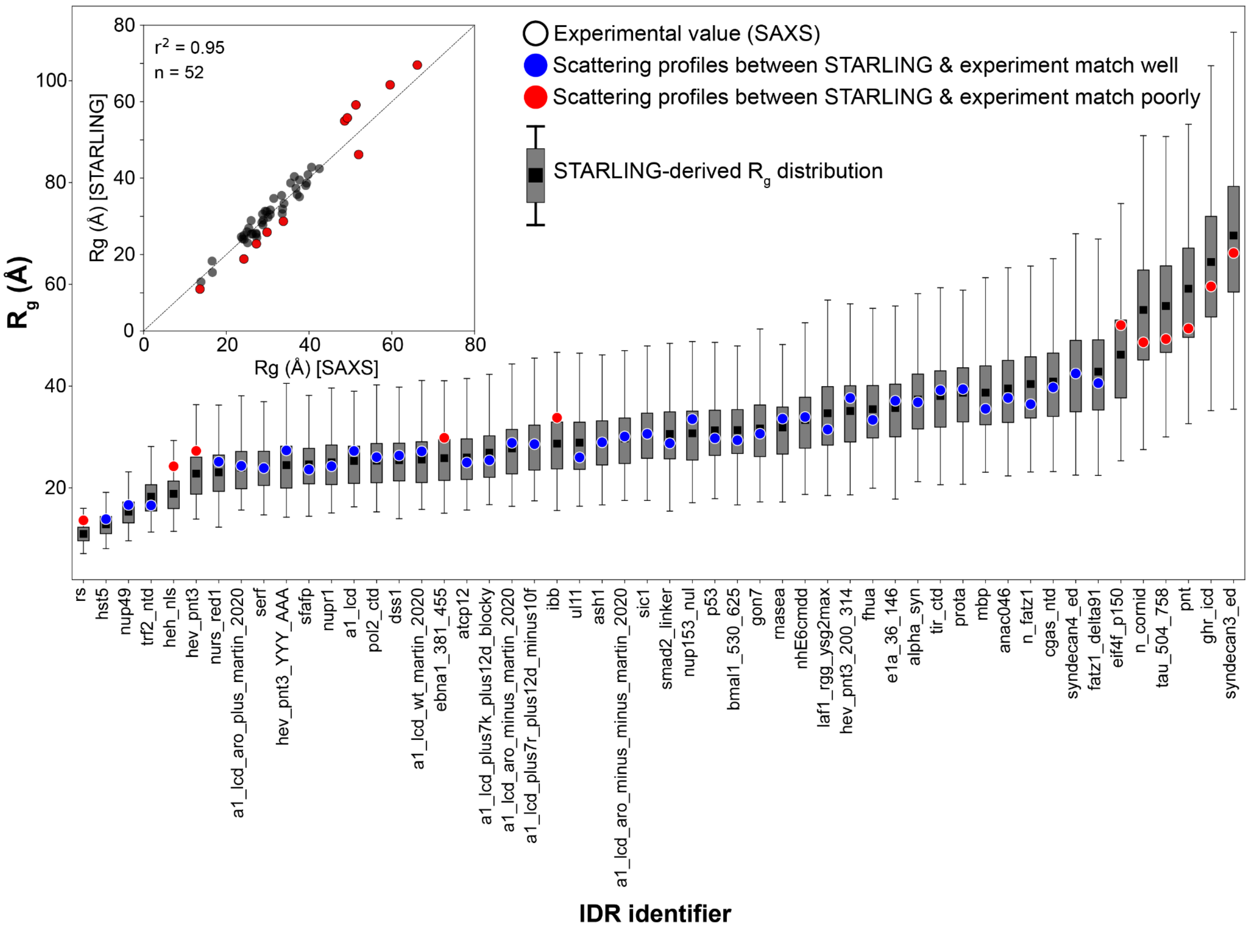


**Figure S14.** Comparison of SAXS-derived radii of gyration vs. STARLING-derived radii of gyration for the same 52 sequences as described in **Fig. S13**. Sequences in which experimental data is represented as a red dot are those where scattering profiles were in sub-optimal agreement with STARLING-derived scattering data—overall, focussing on this high-quality dataset (each of which we re-analyzed to obtain $R_g$ values using the Molecular Form Factor (MFF) of Riback et al.[6] we find even stronger agreement with STARLING-derived predictions (inset).

**A.**

```
>sGrich
GSGSCSGGYGSERGGGYGSERGGGYGSERGGGYGSERGGGYGSQRSGGGYGGSQRSSYGSGSCTLGPR
>dKh
GSGSCAMGGGPGPGTDFTSDQADFPDTLFQEFEPPAPRPGLAGGRPGDAALLSAAYGRRRLLCTLGPR
>dGrich
GSGSCGPRTGLEGAGMAGGSGQQKRVFDGQSGPQDLGEAYRPLNHDGGDGGNRYSVIDRIQECTLGPR
>sPTBP
GSGSCRPDLPSGDSQPSLDQTMAAAFGLSVPNVHGALAPLAIPSAAAAAAAAGRIAIPGLAGCTLGPR
>sCh
GSGSCKKAQPEMNDKDDNESGNEDAEENHDDEEDENEEEDRQVDQASKNKESKRKAQNKREDCTLGPR
>sNrich
GSGSCLDQEDNNGPLLIKTANNLIQNNSNMLPLNALHNAPPMHLNEGGISNMRVNDSLPSNTCTLGPR
>sKh
GSGSCKPRRLSKLRRSKKPADEENNAASQDPTVEATQERGQASEDPENAANNAKQAKPTSDDCTLGPR
>dCh_plus
GSGSCQTPLKRIKVKTPGKSGAAAREGSVVSGTDGPTQTGKPERRKRLNPPKDKLIDMDDADCTLGPR
>dCh_minus
GSGSCMGLPTGMEEKEEGTDESEQKPVVQTPAQPDDSAEVDSAALDQAESAKQQGPILTKHGCTLGPR
>sKl
GSGSCAPEGVFKLPAPPKEIKKSEKSEKSSDSSNDKKEVESTTKTAATTTTTKKGTDNNTQFCTLGPR
>sNh_plus
GSGSCPEEIETRKKDRKNRREMLKRTSALQPAAPKPTHKKPVPKRNVGAERKSTINEDLLPPCTLGPR
>dArich
GSGSCLPTGAEGRDSSKGEDSAEETEAKPAVVAPAPVVEAVSTPSAAFPSDATAEQGPILTKCTLGPR
>dErich
GSGSCEGEMFGVMCEENSVVLSVEQPAELKEVADVSPPPTTRNHTIEMKPPLSAQQSECNPVCTLGPR
>dTRBP
GSGSCLEPALEDSSSFSPLDSSLPEDIPVFTAAAAATPVPSVVLTRSPPMELQPPVSPQQSECTLGPR
>dKl
GSGSCLPTGMDQKTTELMKVDEPVSTQETLPPVIKMEPEPVPITETPTDENARQQGPILTKHCTLGPR
>sNh_minus
GSGSCKKKLQDTPSEPMEKDPAEPETVPDGETPEDENPTEEGADNSSAKMEEEEEEEEEEEECTLGPR
```

**B.**

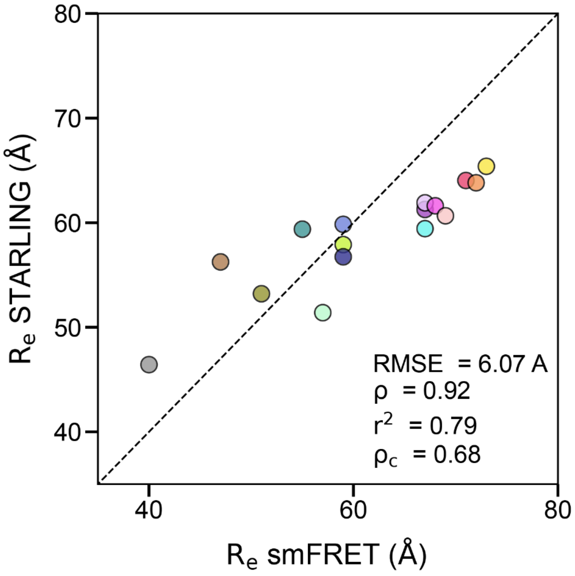


**Fig. S15** Sequences and overall correlation obtained for sequences investigated by smFRET by [7]. **(A)** The amino acid sequence of the sequences investigated. **(B)** Correlation between STARLING-derived end-to-end distances and smFRET-derived end-to-end distances. Agreement between STARLING and experiment here is assessed in various ways, including root mean squared error (RMSE), Spearman correlation coefficient (ρ), the square of the Pearson correlation coefficient ($r^2$), and the concordance correlation coefficient ($\rho_c$). We note that STARLING appears to underestimate end-to-end distances for more expanded chains here, although we do not take dyes into explicit consideration, and it is possible that for the expanded negatively charged sequence, a small amount of (negatively-charged) dye:protein repulsions could account for the slight underestimation here.

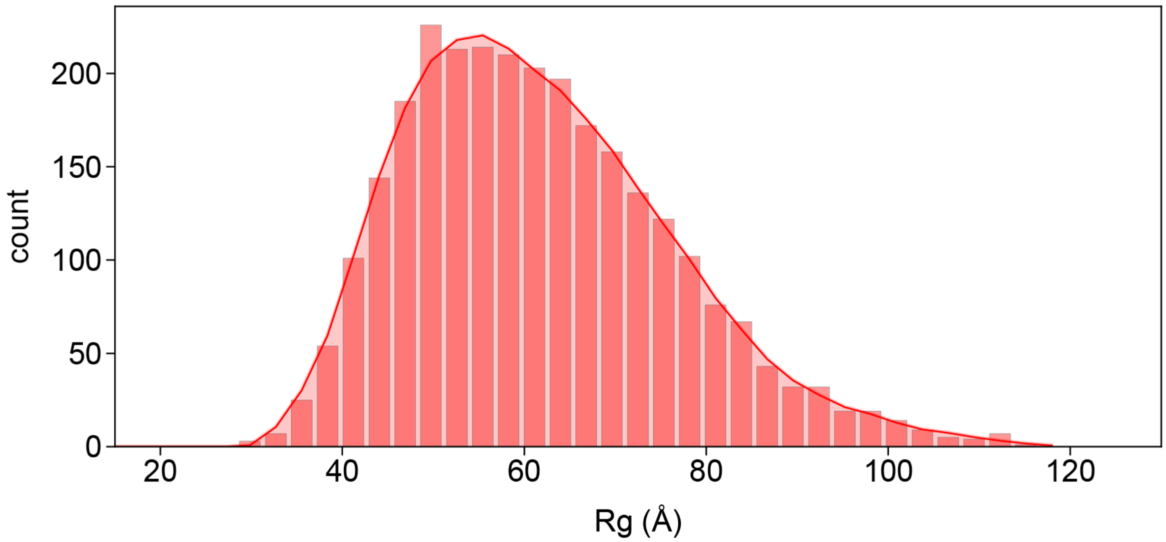


**Fig. S16** $R_g$ distribution for cMyc[1-361] obtained from STARLING ensembles. We predict an average Rg of ~60 Å. We note this value may be an over-estimate given cMyc[1-361] is predicted to contain several regions of transient secondary structure, which may reduce the $R_g$, such that we suggest a value closer to 55 Å may be a more accurate reflection on the expected dimensions for monomeric cMyc[1-361].

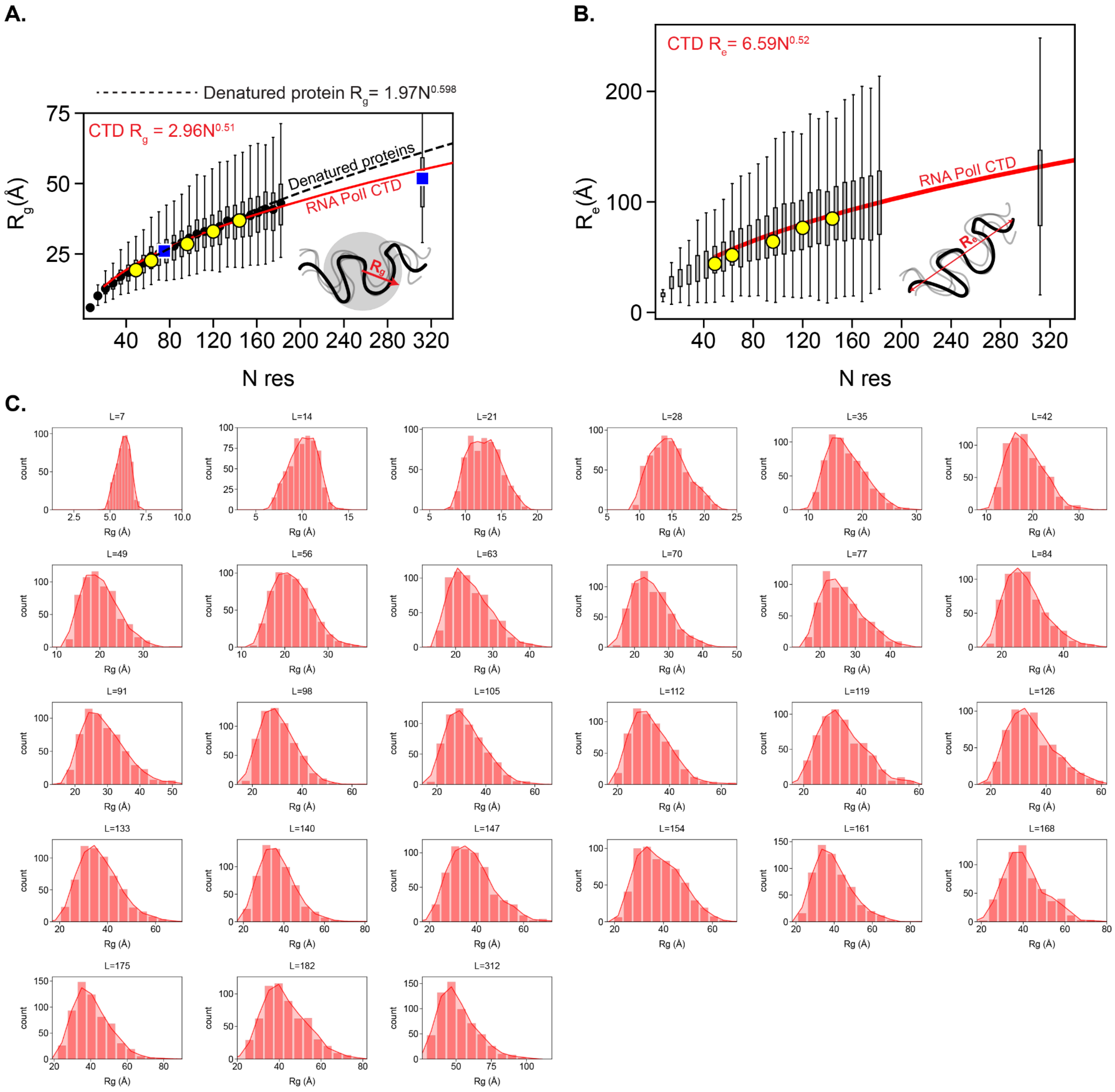


**Fig. S17** Conformational behavior of RNA polymerase II CTD was examined using STARLING ensembles **(A)**. The same figure is shown for **Fig. 4D**, with an additional polymer scaling line here derived from fully denatured proteins measured by Kohn et al., also superimposed (black dashed line)[8]. This is shown because SAXS data on the 83-residue construct of the CTD measured by Gibbs *et al*. revealed a radius of gyration comparable to a fully denatured protein of the same number of amino acids (~28 Å in both cases)[9]. This observation is perhaps unexpected, given that fully denatured proteins are considered to engage in minimal intra (or inter) molecular interactions due to the “good solvent” provided by high denaturant concentrations. Yet, paradoxically, much work has shown that the RNA Pol2 CTD can phase separate and engage in heterogeneous non-random intramolecular contacts[10–13]. These results appear to resolve this apparent discrepancy; the larger dimensions of the 83-residue fragment originate primarily through the larger prefactor value in the scaling relationship ($B_0$ = 2.96 for CTD vs. 1.97 for denatured proteins), presumably driven by the stiffness of the proline residues and – perhaps to a lesser extent – the excluded volume of the tyrosine residue. As such, the CTD does indeed behave in a manner that involves extensive intramolecular interactions, but the overall global dimensions are dominated by the large prefactor (a convolution of chain persistence length and monomer excluded volume). We suggest the presence of proline residues – beyond serving as CKD recognition sites for phosphorylation – serves a biophysical role, enabling such a tyrosine-rich sequence to remain highly expanded and be primed for intermolecular interactions with a cost of cofactors during search, initiation, and elongation. **(B)** The analogous figure is shown in Fig. 4D, but the end-to-end distance is shown instead of the radius of gyration. **(C)** Histograms for $R_g$ values across all CTD lengths.

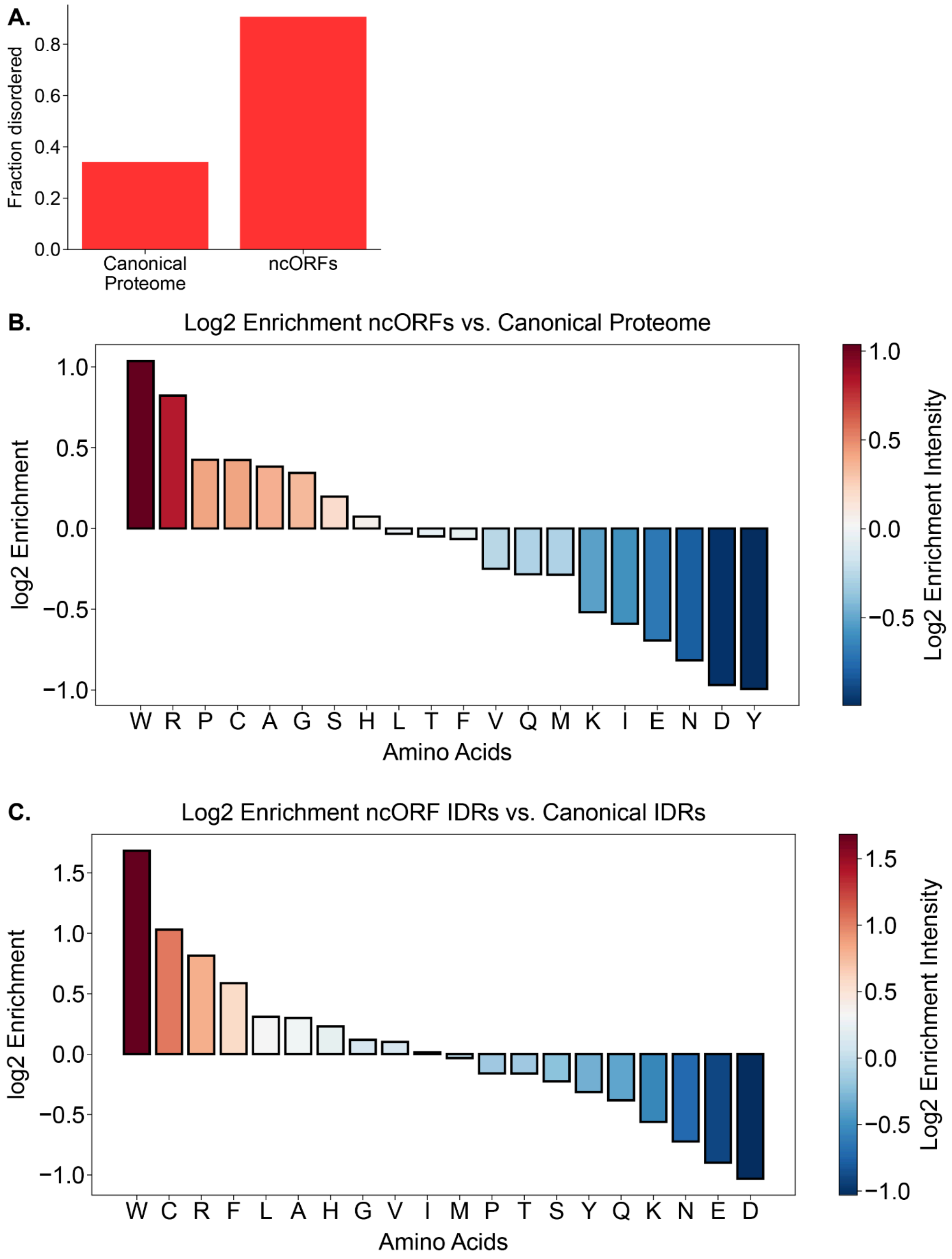


**Fig. S18. (A)** Microproteins are predicted to be highly disordered. The comparison here is the fraction of amino acids in the canonical human proteome found in disordered regions vs. the fraction of amino acids in the Deutsch *et al.* micro-proteome found in disordered regions[14]. The disorder was predicted using metapredict V3[15]. **(B)** Statistical enrichment for each of the 20 amino acids comparing frequency observed across all canonical proteins vs. frequency observed in microproteins. We excluded initiator methionine for both datasets, given the proportional impact of such a bias disproportionally influences shorter vs. longer sequences. As observed previously for metazoan microproteins, we find enrichment for tryptophan and arginine and depletion acid residues[16,17]. **(C)** To establish whether the enrichment observed here reflects something specific to microproteins or the enhanced fraction of disorder, we also compared enrichment in microprotein-derived IDRs vs. canonical-proteome-derived IDRs. This revealed an even stronger signature: microprotein IDRs are *highly* enriched for tryptophan, cysteine, arginine, and phenylalanine. In contrast, they are highly depleted for lysine, asparagine, glutamic acid, and aspartic acid.

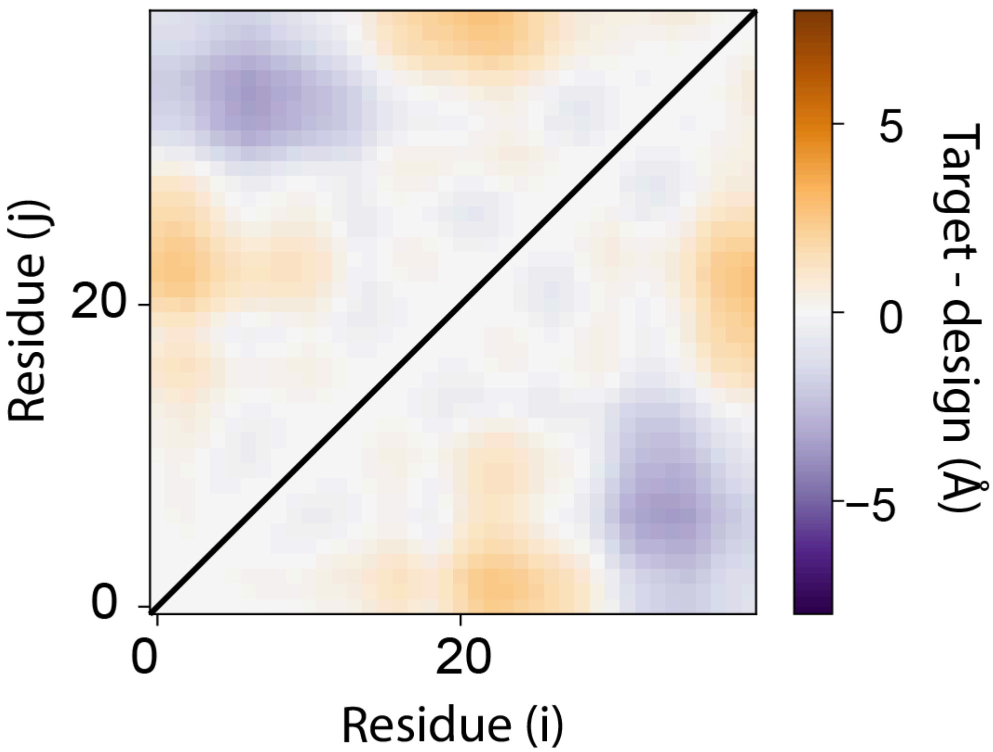


**Fig. S19** Symmetric inter-residue distance matrix showing target designed sequence subtracted from the target sequence, yielding a difference map. All inter-residue errors are < 4 Å, with an RMSE error of 1.3 Å.

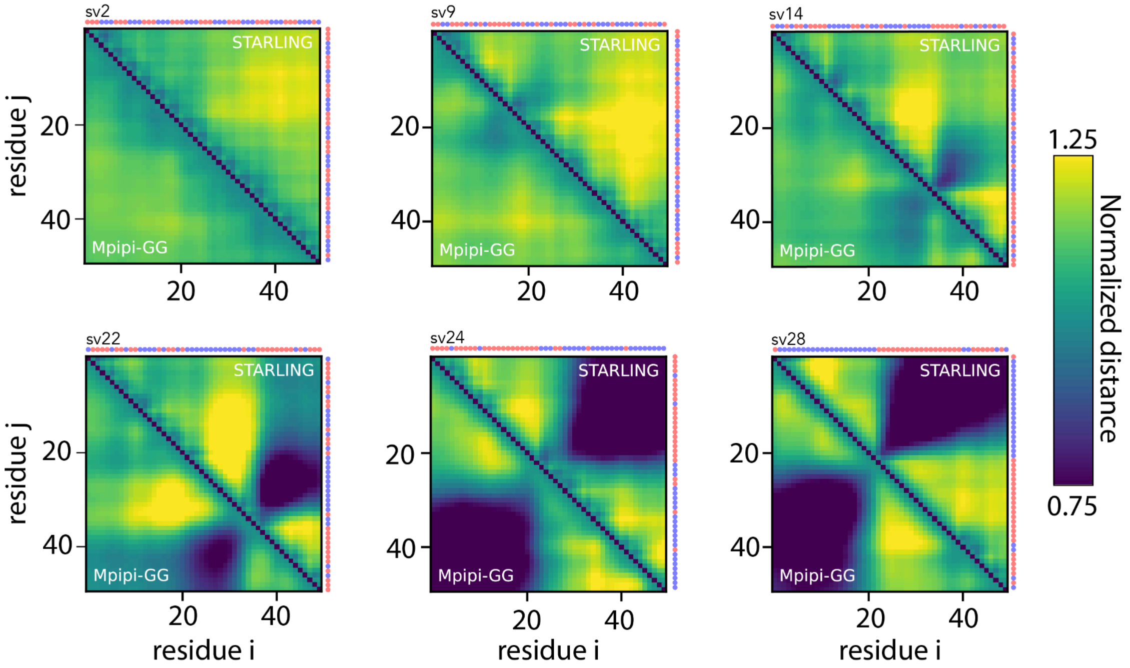


**Fig. S20 Das-Kappa sequences in STARLING vs. Mpipi-GG.** We computed normalized distance maps for several of the iconic "Das sequence"[20], revealing that STARLING ensembles offer good reproduction of the inter-residue distances.

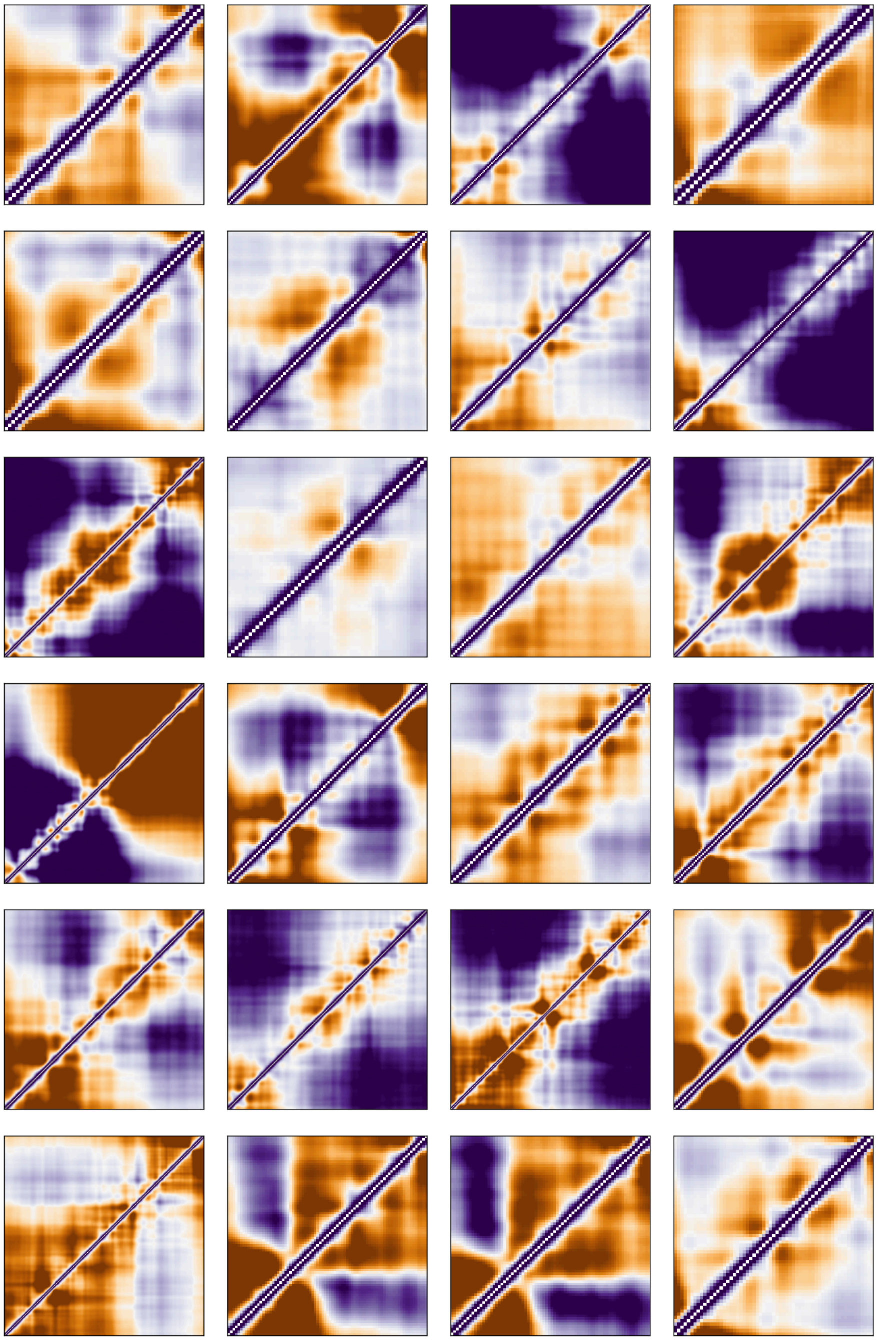

**Fig. S21 Example of distance maps from DisProt**

We examined normalized distance maps from a set of more compact ensembles identified from our DisProt sequences, revealing a wealth of sequence-encoded complexity in the underlying conformational behavior.

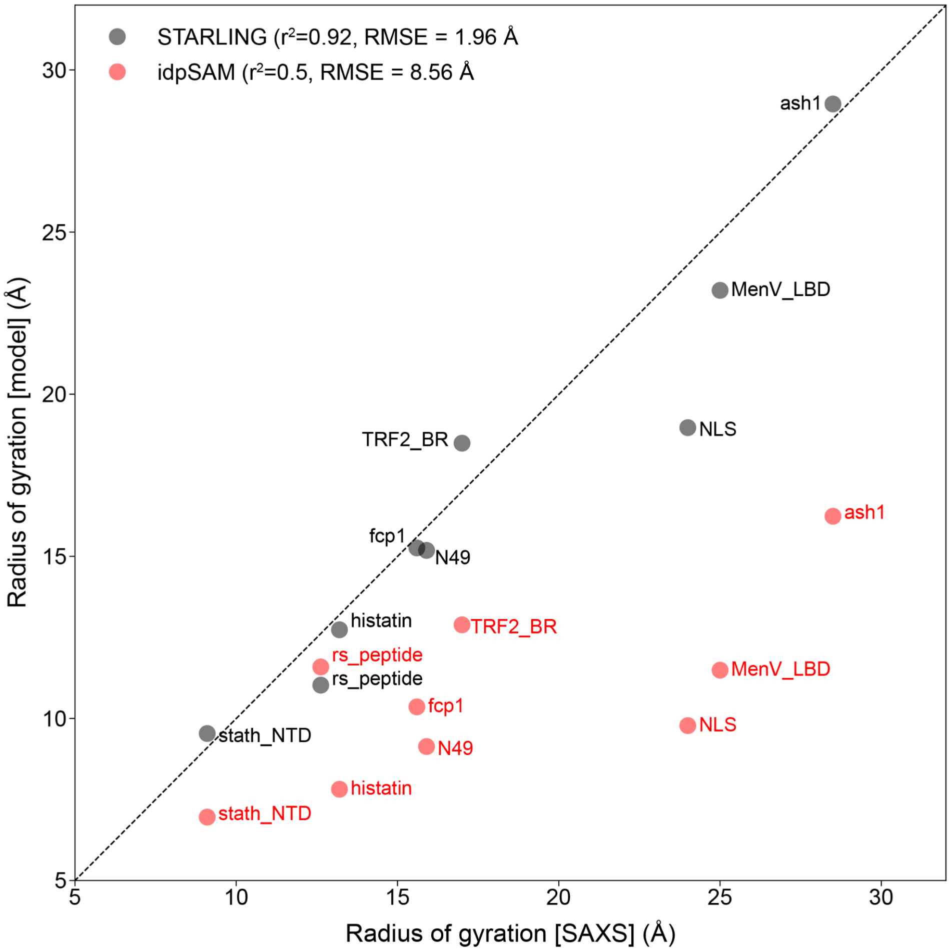


**Fig. S22**. **Comparison of idpSAM-derived ensembles vs. STARLING compared to experimental data.** Focussing on sequences less than 60 amino acids in length (with the exception of Ash1, discussed below), we compared $R_g$ values obtained from idpSAM ensembles vs. $R_g$ values from STARLING ensembles. In both cases, 1000 conformers were used (deferring to the idpSAM defaults). idpSAM offers limited predictive power. We note that the Analytical Flory Random Coil (AFRC) – a null model in which IDR sequence chemistry does not influence the overall dimensions and instead IDR dimensions depend solely on chain length and (weakly) the intrinsic dimensions of each amino acid – obtains an $r^2$ of 0.9 and an RMSE of 3.58 Å[18]. We included Ash1 (83 amino acids) because while it is larger than the ~60 amino acid cutoff used for training data in idpSAM, prior work has shown SAXS data for Ash1 is well reproduced by CAMPARI/ABSINTH simulations[19]. However, despite being trained on CAMPARI/ABSINTH simulations, idpSAM was unfortunately unable to recapitulate this behavior.

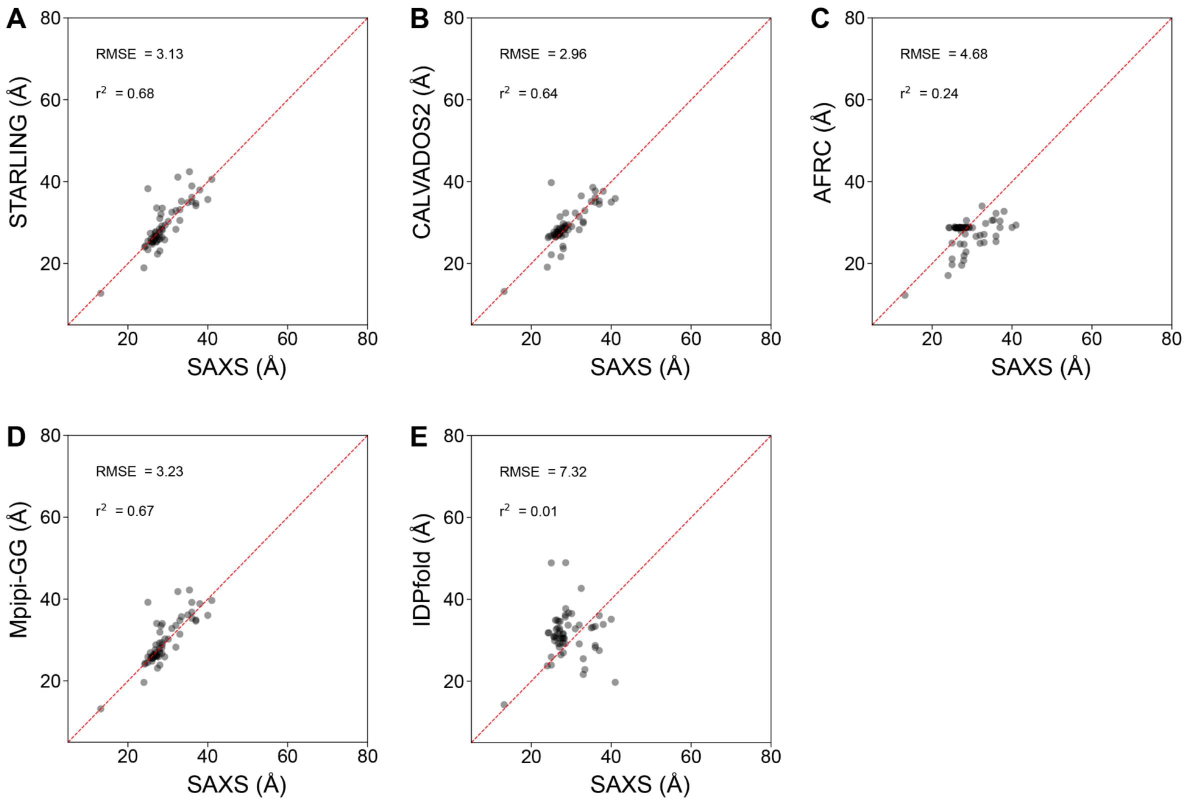


**Fig. S23. Comparison of IDPFold-derived radii of gyration vs. other approaches compared to radii of gyration derived from experimental SAXS data. (A)** Radii of gyration for STARLING vs. SAXS for 59 sequences where IDPFold ensembles were available at the time of submission. **(B)** Radii of gyration for CALVADOS2 simulations vs. SAXS for 59 sequences where IDPFold ensembles were available at the time of submission. **(C)** Radii of gyration for AFRC vs. SAXS for 59 sequences where IDPFold ensembles were available at the time of submission. **(D)** Radii of gyration for Mpipi-GG simulations vs. SAXS for 59 sequences where IDPFold ensembles were available at the time of submission. **(E)** Radii of gyration for IDPFold vs. SAXS for 59 sequences where IDPFold ensembles were available at the time of submission.

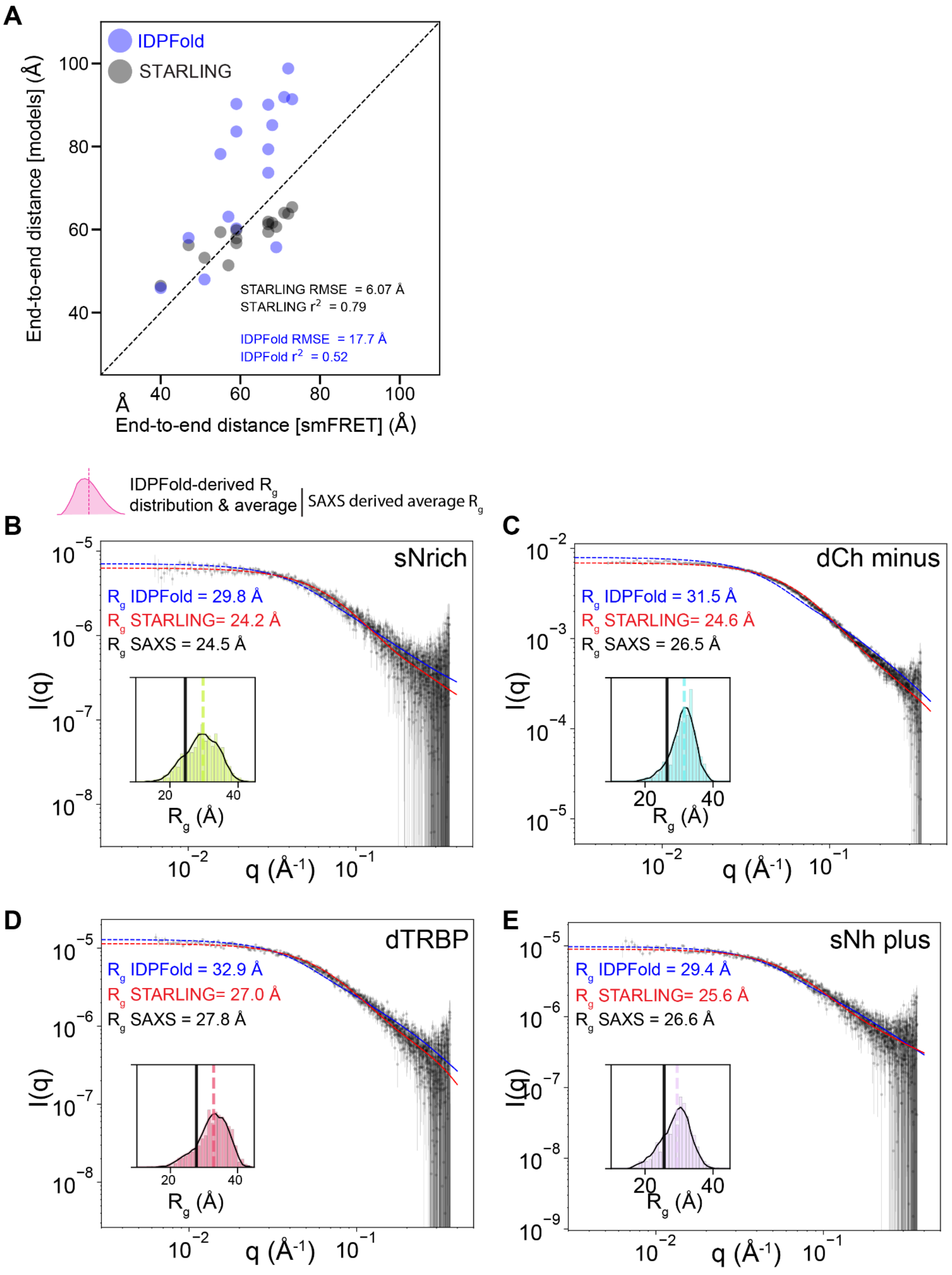


**Fig. S24. Comparison of IDPFold-derived ensembles vs. STARLING compared to single-molecule FRET data. (A)** Correlation between experimentally-derived end-to-end distance (x-axis) and STARLING (black) or IDPFold (Blue) derived end-to-end distances. **(B)** Comparison of SAXS scattering profile for the sNrich ensemble derived from IDPFold ensembles (red) vs. experiment (black). Inferred radii of gyration from IDPFold, STARLING, and SAXS are provided. **(C)** Comparison of SAXS scattering profile for the dCh minus ensemble derived from IDPFold ensembles (red) vs. experiment (black). Inferred radii of gyration from IDPFold, STARLING, and SAXS are provided. **(D)** Comparison of SAXS scattering profile for the dTRBP ensemble derived from IDPFold ensembles (red) vs. experiment (black). Inferred radii of gyration from IDPFold, STARLING, and SAXS are provided. **(B)** Comparison of SAXS scattering profile for the sNh plus ensemble derived from IDPFold ensembles (red) vs. experiment (black). Inferred radii of gyration from IDPFold, STARLING, and SAXS are provided.

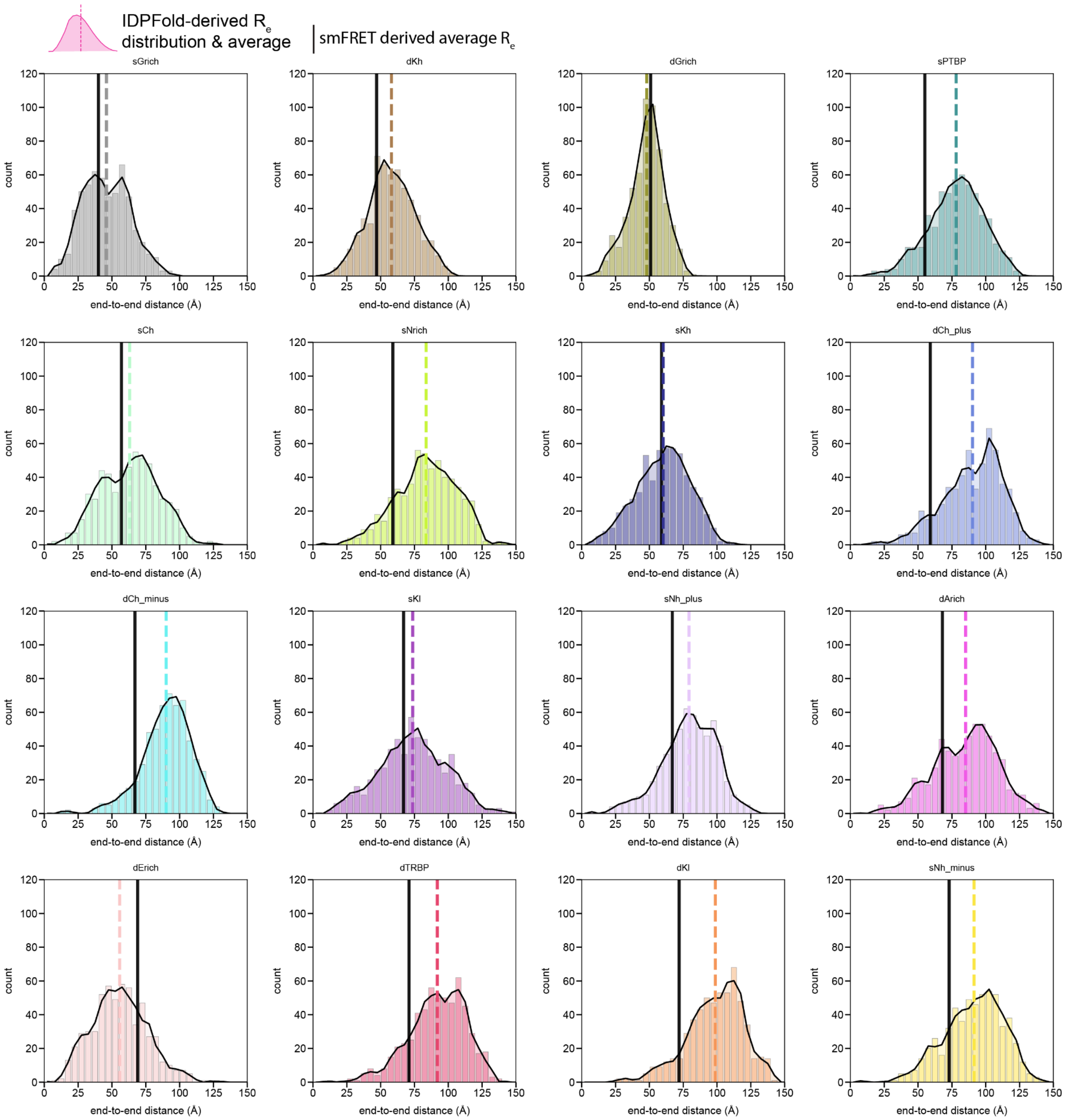


**Fig. S25**. **Comparison of IDPFold-derived ensembles vs. STARLING compared to smFRET data.** Distributions from IDPFold ensembles (colored histograms) with average values (colored dashed lines) for end-to-end distance vs. experimentally derived end-to-end distance (black line).

16. Martinez, T. F., Lyons-Abbott, S., Bookout, A. L., De Souza, E. V., Donaldson, C., Vaughan, J. M., Lau, C., Abramov, A., Baquero, A. F., Baquero, K., Friedrich, D., Huard, J., Davis, R., Kim, B., Koch, T., Mercer, A. J., Misquith, A., Murray, S. A., Perry, S., Pino, L. K., Sanford, C., Simon, A., Zhang, Y., Zipp, G., Bizarro, C. V., Shokhirev, M. N., Whittle, A. J., Searle, B. C., MacCoss, M. J., Saghatelian, A. & Barnes, C. A. Profiling mouse brown and white adipocytes to identify metabolically relevant small ORFs and functional microproteins. *Cell Metab.* **35,** 166–183.e11

(2023).

17. Martinez, T. F., Chu, Q., Donaldson, C., Tan, D., Shokhirev, M. N. & Saghatelian, A. Accurate annotation of human protein-coding small open reading frames. *Nat. Chem. Biol.* **16,** 458–468 (2020).

18. Alston, J. J., Ginell, G. M., Soranno, A. & Holehouse, A. S. The Analytical Flory Random Coil Is a Simple-to-Use Reference Model for Unfolded and Disordered Proteins. *J. Phys. Chem. B* **127,** 4746–4760 (2023).

19. Martin, E. W., Holehouse, A. S., Grace, C. R., Hughes, A., Pappu, R. V. & Mittag, T. Sequence Determinants of the Conformational Properties of an Intrinsically Disordered Protein Prior to and upon Multisite Phosphorylation. *J. Am. Chem. Soc.* **138,** 15323–15335 (2016).

20. Das, R. K. & Pappu, R. V. Conformations of intrinsically disordered proteins are influenced by linear sequence distributions of oppositely charged residues. *Proc. Natl. Acad. Sci. U. S. A.* **110,** 13392–13397 (2013).
